# Characterization of early and late events of adherens junction assembly

**DOI:** 10.1101/2024.03.04.583373

**Authors:** Regina B. Troyanovsky, Indrajyoti Indra, Sergey M. Troyanovsky

**Author notes:** Please address all correspondence to: Dr. Sergey Troyanovsky Department of Dermatology Northwestern University The Feinberg School of Medicine, 303 E. Chicago Ave Chicago, IL 60611.

## Abstract

Cadherins are transmembrane adhesion receptors. Cadherin ectodomains form adhesive 2D clusters through cooperative *trans* and *cis* interactions, whereas its intracellular region interacts with specific cytosolic proteins, termed catenins, to anchor the cadherin-catenin complex (CCC) to the actin cytoskeleton. How these two types of interactions are coordinated in the formation of specialized cell-cell adhesions, adherens junctions (AJ), remains unclear. We focus here on the role of the actin-binding domain of α-catenin (αABD) by showing that the interaction of αABD with actin generates actin-bound CCC oligomers (CCC/actin strands) incorporating up to six CCCs. The strands are primarily formed on the actin-rich cell protrusions. Once in cell-cell interface, the strands become involved in cadherin ectodomain clustering. Such combination of the extracellular and intracellular oligomerizations gives rise to the composite oligomers, *trans* CCC/actin clusters. To mature, these clusters then rearrange their actin filaments using several redundant pathways, two of which are characterized here: one depends on the α-catenin-associated protein, vinculin and the second one depends on the unstructured C-terminus of αABD. Thus, AJ assembly proceeds through spontaneous formation of *trans* CCC/actin clusters and their successive reorganization.

## Introduction

Adherens Junctions (AJs), evolutionary the oldest type of cell-cell adhesions, establish strong but flexible contacts between cells [1–5]. This remarkable property allows cells to stay connected during tissue morphogenesis and is based on continuous disassembly and reassembly of numerous actin-bound adhesion clusters of classic cadherins (e.g., E-cadherin in epithelia). While the structures of cadherin adhesive (or *trans*) bonds in these clusters and the bonds connecting the clusters to actin filaments are known at atomic detail [6–9], coordination of these two sets of interactions during cadherin cluster lifetime is far from being understood. Elucidation of cadherin clustering is critical for our understanding of the adhesion defects that are known to associate with many human diseases [10, 11].

The structural unit of cadherin clusters is the cadherin-catenin complex (CCC), in which the intracellular region of cadherin interacts with two cytosolic proteins, p120-catenin and β-catenin, the latter of which interacts with the actin-binding protein, α-catenin. Cadherin clusters greatly reinforce the strength of an intrinsically weak adhesive *trans* bond [12]. The clusters can be self-assembled through cooperative *trans* and *cis* interactions of the cadherin extracellular region (ectodomain) without any contribution from cytosolic proteins [6, 13–18]. However, the ectodomain-assembled clusters (designated as *trans* E-clusters below) are still too weak to maintain sufficient cell-cell adhesion strength [19, 20]. The stability of *trans* E-clusters is upregulated by coupling them to the actin cytoskeleton through α-catenin that not only provides a structural foundation for the clusters but also reinforces both cadherin *trans* bonds and the bonds between α-catenin and actin filaments via coupling them to the actomyosin tensile forces [3, 18, 21, 22]. It has also been shown that α-catenin binding to F-actin preferentially occurs in the vicinity of already bound α-catenin [23]. Such remarkable cooperativity of α-catenin binding to actin was proposed to generate CCC clusters independently of the *trans* E-cluster formation [24]. Whether this actin-based α-catenin clustering plays any role in AJs and if it does, how intracellular and extracellular clustering are coordinated remain unexplored.

A key actin-binding component of α-catenin is its C-terminal domain, termed here αABD (see Fig. 1 for detail). Cryo-EM modeling of the αABD-actin complex suggests that αABD simultaneously interacts with the actin filament and with two neighboring actin-bound αABDs, thereby forming an actin-bound linear αABD oligomer [8, 9, 25]. In vitro binding assays also suggest that this binding mode, called the strong binding state, is facilitated by a pulling force applied on the initially weak αABD-actin bond [22, 26–28]. Taken together, this data could be interpreted that the strong αABD-actin interaction follows the *trans* dimerization of the cadherin ectodomain, which allows an application of force. However, several observations made in cell culture appear inconsistent with the role of force and/or *trans* interactions in the weak-to-strong αABD-actin bond conversion. For example, the actin bound CCC clusters were detected on the contact-free surface of the Drosophila cells [16]. The actomyosin-dependent stretching of E-cadherin had been detected outside of cell-cell contacts [29]. The anti-actomyosin drugs have not been shown to abolish AJs, but only change their appearance [30–33]. This data suggests that CCC may associate with actin in both an adhesion- and force-independent manner. These inconsistencies in binding assays and in cell culture necessitate the need for new tools in order to be able to directly monitor the strong actin binding state of αABD in living cells.

**Figure 1.**
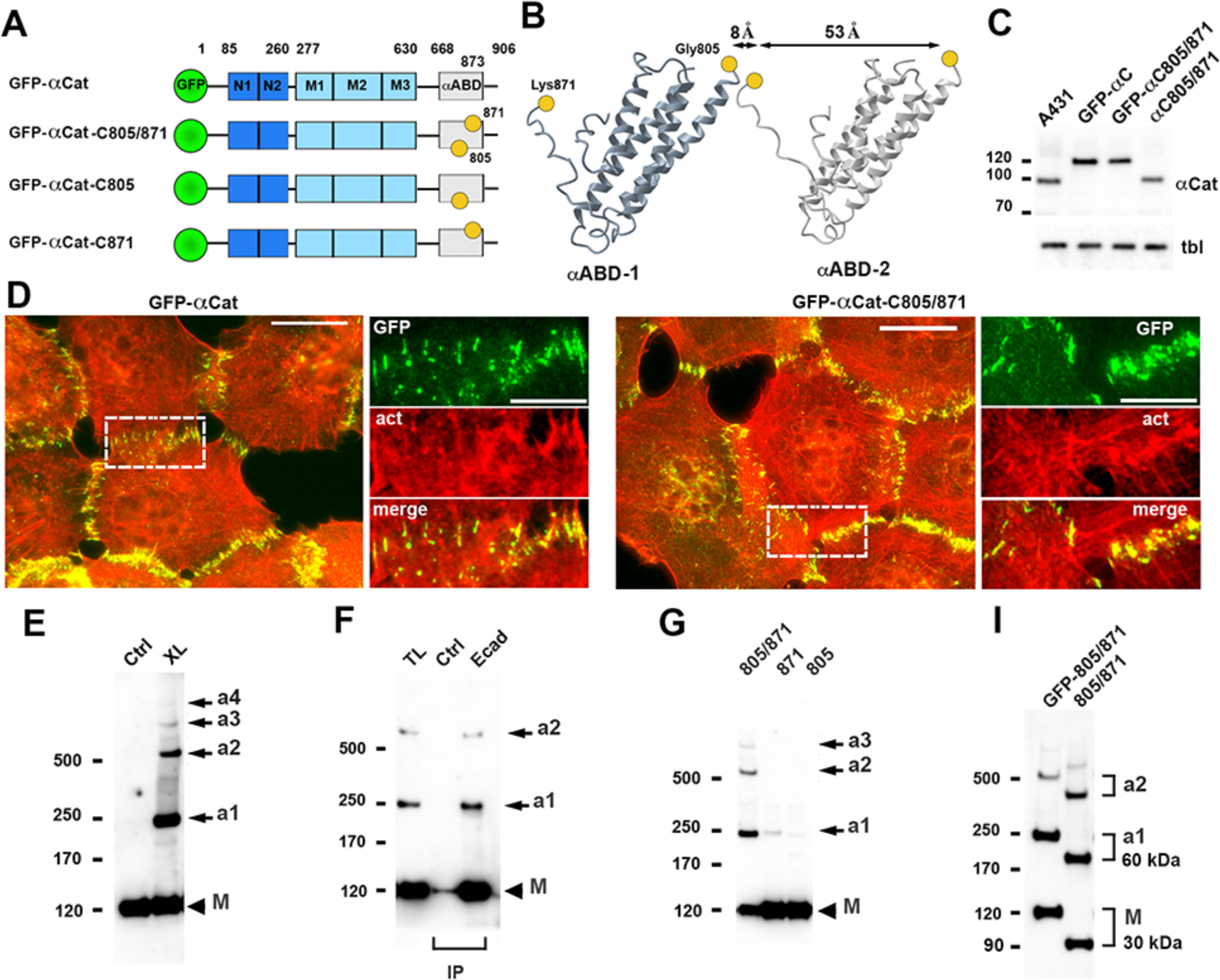
Detection of α-catenin oligomers. (**A**) Diagram of the α-catenin mutants used for targeted cross-linking: GFP (green), N domain (dark blue) comprising of N1 and N2 subdomains; M domain (light blue) comprising of M1, M2, and M3 subdomains, and actin-binding domain, αABD (grey). The unstructured regions, including the αABD tail (residues 873-906), are shown as solid lines. The borders between domains are indicated by the numbers of corresponding residues. Cysteine substitutions, G805C (805), and K871C (871) are shown as yellow spheres. (**B**) Ribbon diagram of two neighboring actin-bound αABD (αABD-1 and αABD-2) according to PDB 6wvt. The cysteine substitutions are shown as in A. Note, that the G805C/K871C mutation creates an ideal cysteine pair for cross-linking of the contiguous protomers. (**C**) Western blot of the total lysates of wt A431 cells (A431), and αCatKO-A431 cells expressing GFP-αCat (GFP-αC), GFP-αCat-C805/871 (GFP-αC805/871), and untagged αCat-C805/871 (αC805/871). The blot was probed for α-catenin (αCat) and for tubulin (tbl) as a loading control. Molecular weight markers (in kDa) are shown on the left. (**D**) Fluorescence microscopy of αCatKO-A431 cells expressing GFP-αCat and GFP-αCat-C805/871 stained for GFP (GFP, green) and for F-actin (act, red). The left micrographs show merged images (Scale bar, 25 μm). The separate images of the representative cell-cell contacts (in dashed boxes) are zoomed on the right (Scale bar, 10 μm). Note that cells of both lines form actin-associated AJs. (**E**) Western blot of total lysates of cells expressing GFP-αCat-C805/871 probed for GFP. Without cross-linking (Ctrl), the mutant migrates as a single band of ∼120kDa (M). Cross-linking (XL) results in formation of adducts (a1-4). (**F**) A lysate of cross-linking cells (TL) was split into two parts and immunoprecipitated (IP) with E-cadherin mAb (Ecad) or without a first antibody (Ctrl). The precipitates were processed as in E. (**G**) Cells expressing GFP-αCat-C805/871 (805/871) or its versions with a single substitution (805 or 871) were cross-linked and processed as in E. (**I**) Cells expressing GFP-αCat-C805/871 mutant (GFP-805/871) or its untagged version (805/871) were cross-linked and analyzed as in E using an α-catenin antibody. Note that the difference in MW of a1 between tagged and untagged mutants is about 60 kDa, twice more that the difference (∼ 30 kDa) between the monomers.

Here we track the strong αABD-actin bonds using a targeted cross-linking approach. Our results present the compelling evidence that the two types of CCC oligomerization, formation of *trans* E-clusters and linear oligomerization of αABD on actin filaments, are independent, but could integrate with one another producing composite oligomers, *trans* CCC/actin clusters. We also show that formation of these clusters is not sufficient for stable adhesion. Additional processes, which include the rearrangement of the cluster-associated actin filaments, are required to assemble the adhesion-competent AJs.

## Results

### Identification of a-catenin oligomers in cells

To analyze the actin-bound CCC clusters, we applied targeted chemical cross-linking, a technique we have used to study cadherin and nectin *trans* interactions [34, 35]. Specifically, using a cryo-EM map of an actin-bound αABD (PDB entry 6wvt), we designed an αABD cysteine mutant, GFP-αCat-C805/871. Cysteine substitutions in this mutant, G805C and K871C, were positioned within two unstructured αABD regions, in the α4/α5 loop and in the portion of αABD C-terminal extension (referred as αABD tail below), respectively (Fig. 1A, B). According to the map, the intramolecular distance between these residues in an αABD is ∼ 50 Å, whereas the intermolecular distance between the same residues from two adjacent actin-bound αABDs is only 9 Å (Fig. 1B). Because of this difference, the cell permeable sulfhydryl-reactive cross-linker, 1,4-bismaleimidobutane (BMB, about 10 Å in size), should preferentially cross-link adjacent actin-bound αABDs. Also, according to the cryo-EM, the mutated residues are not involved in interactions with F-actin. In agreement with this, the GFP-αCat-C805/871 mutant, similar to the intact α-catenin, GFP-αCat, rescued the formation of actin-associated AJs (Fig. 1C, D) upon expression in α-catenin knockout A431 cells (αCatKO-A431).

Western blotting showed that treatment of the mutant expressing cells with a low BMB concentration (40 μM) resulted in the formation of a ladder of high molecular weight adducts (Fig. 1E) whereby the lowest adduct in the ladder (a1 in Fig. 1E) exactly matched the size of the GFP-αCat-C805/871 dimer (∼ 240 kD). Other major adducts higher than 350 kDa (a2 and a3, observed in all performed experiments, and a4, observed occasionally) most likely incorporated three or more cross-linked α-catenin molecules. We then verified that the adducts were derived from CCC, but not from α-catenin homodimers, which are known to form in the cytosol and can interact with F-actin [36, 37]. To this end, we precipitated CCC using an anti-E-cadherin antibody from the lysates of cross-linked GFP-αCat-C805/871-expressing cells. Western blotting of the precipitates clearly showed that the adducts associated with E-cadherin (Fig. 1F).

To exclude the possibility that the adducts were formed by cross-linking of the α-catenin mutant to irrelevant proteins, we constructed two additional α-catenin mutants, GFP-αCat-C805 and GFP-αCat-C871, which harbored only G805C or only K871C substitutions (Fig. 1A). Only barely visible α-catenin adducts could be seen after cross-linking of the cells expressing either of these two mutants (Fig. 1G). This observation confirmed that the adducts were formed through cross-linking of adjacent α-catenin molecules via new cysteines, C805 and C871. In another experiment, we introduced the G805C and K871C substitutions into untagged α-catenin (αCat-C805/871). The deletion of GFP tag decreased the molecular weight of the mutant by ∼ 30 kDa (from ∼120 kDa to 90 kDa). As expected for its dimer organization, the molecular weight of the 240 kDa adduct was decreased by 60 kDa (Fig. 1I). The sizes of other adducts also decreased by more than just by 30 kDa thereby verifying that they all were α-catenin oligomers. Importantly, this experiment also showed that the GFP tag had no influence on the mutant cross-linking.

### Actin filaments are required for a-catenin oligomerization

We next verified that the oligomers we detected required actin filaments for their formation. We introduced a new mutation, I792A, which was shown to decrease α-catenin binding to F-actin [24], into the GFP-αCat-C805/871 mutant. As expected, a resulting triple mutant, GFP-αCatI792A-C805/871, was unable to form AJs in αCatKO-A431 cells (Fig. 2A) and generated only a negligible amount of adducts (Fig. 2B). The role of F-actin in formation of α-catenin oligomers was further substantiated by a strong reduction of the adducts in cells treated with latrunculin B (LnB), which prevents actin polymerization (Fig. 2C). Some amounts of adducts remaining in the treated cells were apparently derived from α-catenin bound to LnB-resistant filaments. Such interactions had been reported [38].

**Figure 2.**
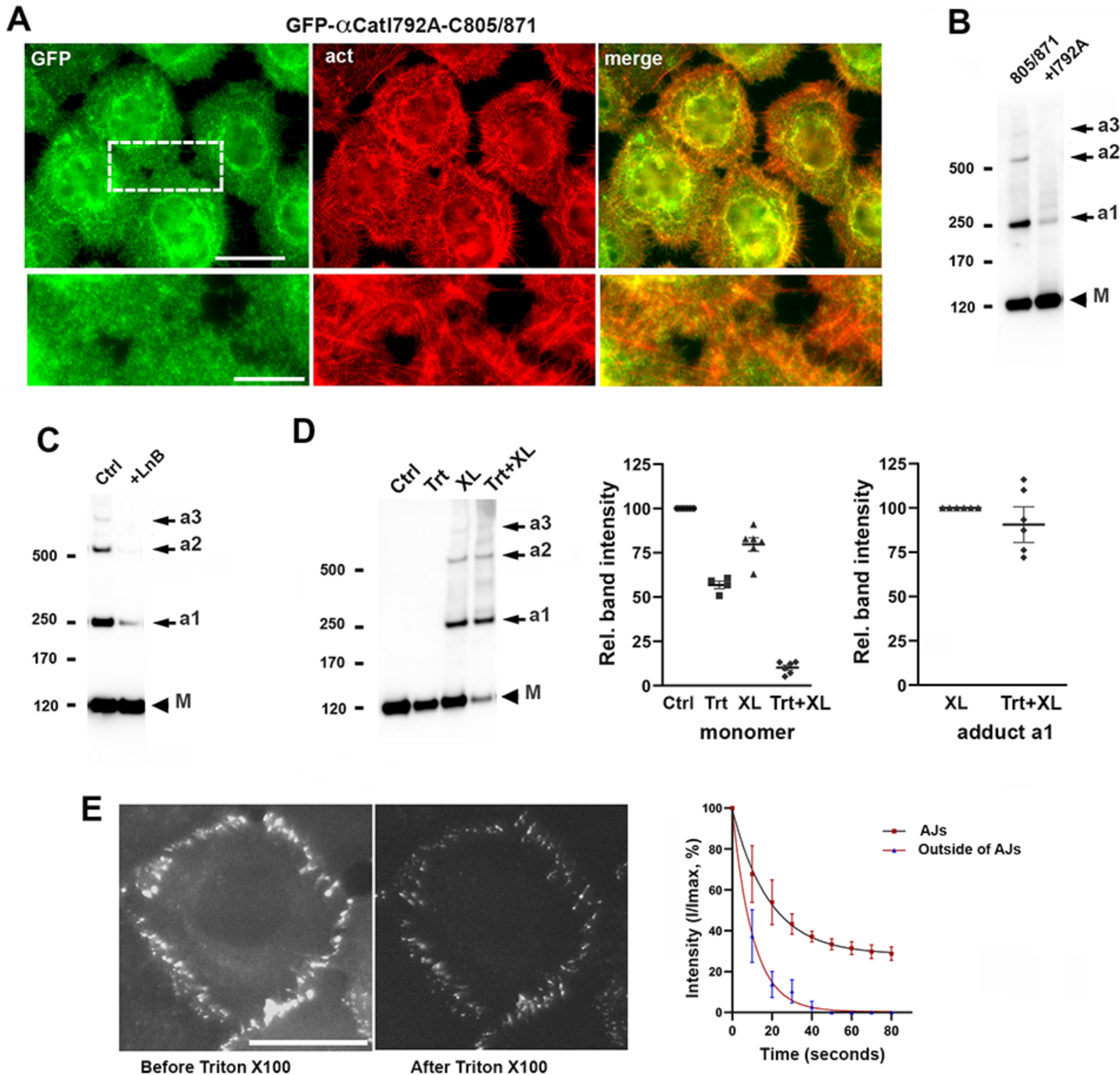
Actin filaments are required for α-catenin oligomerization. (**A**) Fluorescence microscopy of αCatKO-A431 cells expressing actin-uncoupled GFP-αCatI792A-C805/871 mutant stained for GFP (GFP, green) and for F-actin (act, red). Scale bar, 25 μm. A representative cell-cell contact (in a dashed box) is magnified on the bottom (Scale bar, 10 μm). The mutant cannot form AJs. (**B**) Cells expressing GFP-αCat-C805/871 (805/871) and its I792A mutant (+I792A) were cross-linked and processed as in Fig. 1E. (**C**) The GFP-αCat-C805/871 cells were cross-linked in standard conditions (Ctrl) and after 30 min with 1 μM Latrunculin B (LnB). (**D**) Left: Western blot probed for GFP of total lysates of GFP-αCat-C805/871 cells from control culture (Ctrl); from parallel culture after 5 min-long extraction with 1% Triton X100 (Trt); after cross-linking of the control culture (XL); and after cross-linking of the Triton X100 extracted culture (Trt+XL). Right: Quantification (based on five independent experiments) of the results shown on the left. Intensities of the GFP-αCat-C805/871 monomer (relative to that in control culture) and the GFP-αCat-C805/871 adduct, a1 (relative to that in nonextracted culture. The medians + SD are indicated by bars. Note that Triton X100 extraction reduced to ∼50% the level of monomers in the control cells and nearly completely in the cross-linked cells. But it only negligibly changed the level of the adduct. (**E**) Time-lapse microscopy of GFP-αCat-C805/871 cells during Triton X100 extraction. Left: The first (before extraction) and the last (after 1.5 min in Triton X100) frames from one of the obtained representative movies. Bar, 25 μM. Right: Kinetics of the GFP-αCat-C805/871 extraction obtained for AJs and for the spots outside. The error bars represent SDs (n=15).

To expand our data that the adducts were generated from the actin-bound pool of GFP-αCat-C805/871, we employed a gentle extraction of the cells with a Triton X100-containing cytoskeleton stabilization buffer. This approach is a widely accepted procedure to determine proteins interacting with the cytoskeleton [39–41]. The results showed that ∼50% of GFP-αCat-C805/871 remained bound to the cytoskeleton after Triton X100 extraction (Fig. 2D). Cross-linking of the parallel cultures of control and extracted cells demonstrated that about 80% of the Triton X100 resistant pool of GFP-αCat-C805/871 was converted into the adducts. Noteworthy, the amounts of the adducts generated in control and in extracted cells were nearly identical (Fig. 2D). Such almost complete conversion of the Triton X100-resistant α-catenin into adducts demonstrated a remarkable efficiency of cross-linking of the actin-bound GFP-αCat-C805/871. It also validated that Triton X100 specifically removed the actin-uncoupled pool of α-catenin, which allowed us to analyze the size of this actin-free pool of α-catenin in AJs using time-lapse microscopy (Fig. 2E). The results showed that a two min-long extraction of the cells decreased the GFP-αCat-C805/871 fluorescence in AJs by ∼60%, while outside of AJs it dropped to nearly background levels. Collectively, these data showed that our cross-linking approach generates oligomeric adducts exclusively from the actin-bound pool of α-catenin.

### Actin-bound pentameric array of CCC is a predominant product of α-catenin binding to actin

The high molecular weights of these adducts had precluded the exact assessment of their sizes and, therefore, the order of α-catenin oligomerization. In addition, high molecular weight proteins may exhibit low transfer efficiency in Western blotting that hinder their detection and quantification. To get more insight into the adduct organization, we designed an approach allowing us to reduce the adduct molecular weight by cleaving the cross-linked αABD from the rest of the α-catenin. To this end, we incorporated a thrombin cleavage site (TCS) into the linker region between αABD and the M domain by changing the sequence LIAGQS (663-668) to LVPRGS (Fig. 3A). This change had no effect on AJ appearance (Fig. 3B) or the pattern of cross-linked adducts (Fig. 3C). Control experiment with anti-GFP immunoprecipitates obtained from the cells expressing the new mutant (GFP-αCatTCS-C805/871) confirmed that thrombin cut off the αABD (Fig. 3D). We then treated the cells that express the mutant with thrombin after they had been cross-linked and permeabilized with Triton X100 (Fig. 3E). Western blotting with an anti-αABD antibody showed that this procedure yielded a ladder of the cross-linked αABD consisting of four major bands of ∼ 60, ∼90, ∼120, and ∼ 150 kDa. These molecular weights exactly corresponded to those of the dimers, trimers, tetramers and pentamers of αABD. As expected, the monomeric form of αABD was hardly detectable in this experiment due to it removal during Triton-X100 extraction (see above). The most remarkable features of this pattern, however, was the nearly complete absence of the oligomers of an order of 6 or greater. Also, it is interesting to note that the oligomers corresponding to tetramers and pentamers consistently showed approximately the same intensities. We next assessed the relative abundance of each of αABD oligomers. Based on four independent experiments, we determined an Oligomer Intensity Index (OII) - the ratio between the intensity of the oligomer n to that of n-1. It showed that the OII for trimers and tetramers varied between the values of 0.4 to 0.5 (Fig. 3E), while the OII for pentamers was much higher, ranging between 0.9 to 1.1. The index dropped to ∼ 0.2 for the hexamers. Such а nonlinear OII distribution can be interpreted as F-actin generating oligomers comprised of two to six CCCs, among which the pentamer is the most dominant species. To verify that the low amounts of hexamers and the absence of higher-order oligomers was not due to the failures in their detections, we constructed a mutant, GFP-TCS-αABD, consisting of GFP, TCS, and a strong actin-binding mutant of αABD, its truncated form (residues 673-906, see Fig. 3A, F) [8, 9, 23, 26]. It had been shown that the truncated αABD produces numerous short-lived actin-associated clusters when expressed in cells [24]. Therefore, we anticipated that GFP-TCS-αABD would generate high amounts of high-order αABD oligomers. Immunofluorescence microscopy of the αCatKO-A431 cells expressing GFP-TCS-αABD verified that the mutant associated with the actin cortex (Fig. 3G). Cross-linking and thrombin treatment of these cells produced αABD oligomers of up to order 8 (∼ 240 kDa). The OII of all these species varied in range of 0.45 to 0.55 (Fig. 3H). Thus, in complete agreement with structural model, the isolated strong-actin binding mutant of αABD forms a large variety of actin-bound oligomers without a preference for the particular species.

**Figure 3.**
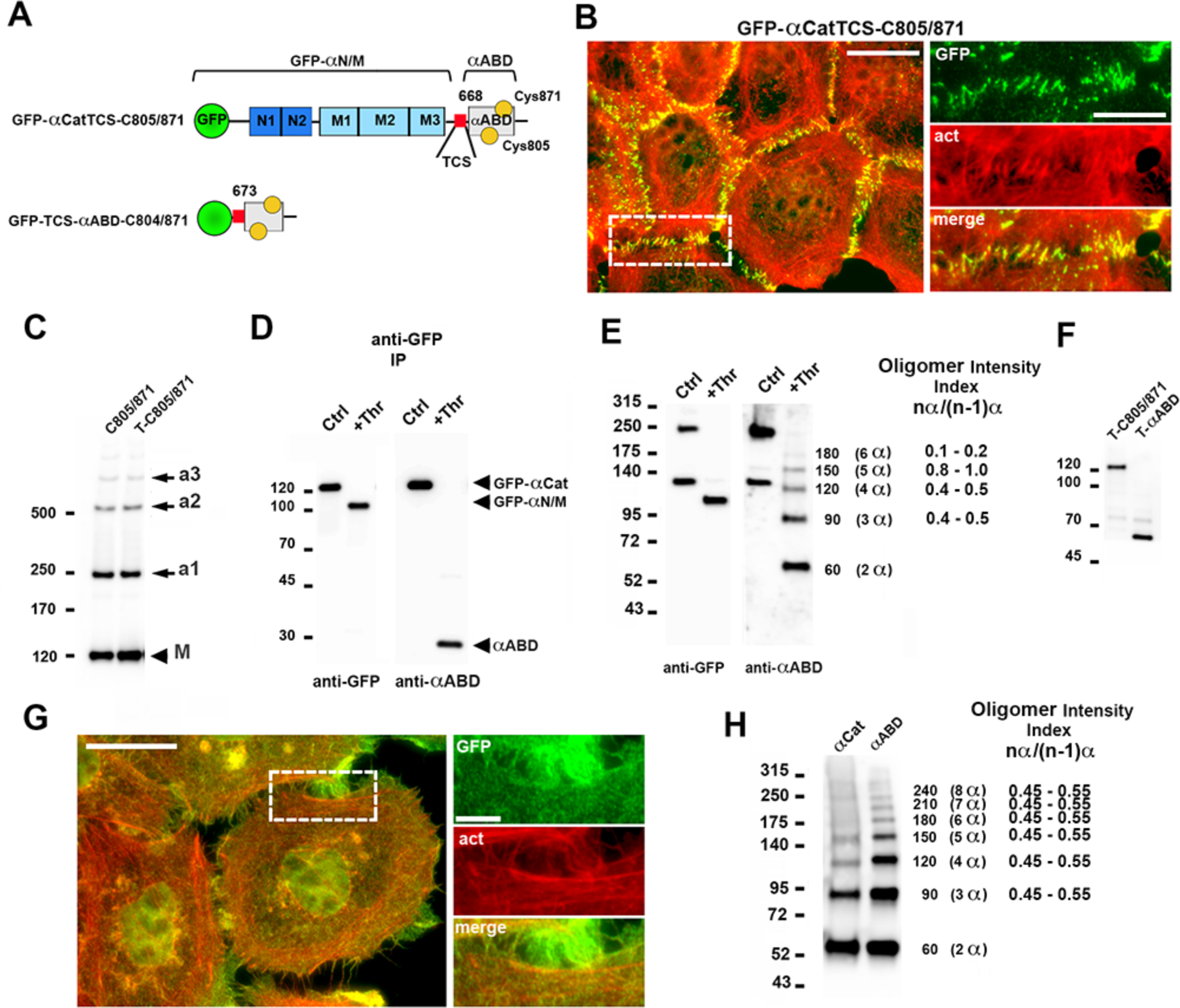
Majority of the actin-bound αABD are pentamers. (**A**) Diagram of the α-catenin mutants used for αABD cleaving. The color code of the α-catenin subdomains is the same as in Figure 1A. The thrombin cleavage site (TCS, red) is positioned in the M-αABD linker that separates the GFP-αCatTCS-C805/871 into two fragments, GFP-αN/M and αABD. TCS is placed between GFP and αABD in GFP-TCS-αABD-C805/871. The αABD in the latter construct starts from the aa 673. (**B**) Fluorescence microscopy of GFP-αCatTCS-C805/871-expressing cells imaged for GFP (GFP, green) and for F-actin (act, red). A representative contact (in a dashed box) taken from the merged image (on the left, Scale bar, 25 μm) is magnified on the right (Scale bar, 12 μm). Note that TCS does not prevent formation of the actin-associated AJs (compare with Fig. 1D). (**C**) Anti-GFP Western blot of total lysates of cross-linked GFP-αCat-C805/871 (C805/871) and GFP-αCatTCS-C805/871 (T-C805/871) cells. Note that both mutants form indistinguishable patterns of adducts. (**D**) Anti-GFP (anti-GFP) or anti-αABD (anti-αABD) Western blotting of the anti-GFP immunoprecipitate of GFP-αCatTCS-C805/871. Before adding the SDS sample buffer, the equal portions of the precipitate were treated for 30 min with or without thrombin (+Thr or Ctrl, correspondingly). Note that thrombin separates the mutant (marked as GFP-αCat) into two fragments, αABD and GFP-αN/M. (**E**) The cells expressing GFP-αCatTCS-C805/871 were permeabilized, treated with (+Thr) or without (Ctrl) thrombin for 30 min, and then separated by SDS-PAGE. Left: Western blots probed by anti-GFP (anti-GFP) or by anti-αABD (anti-αABD). Note, αABD staining showed four bright bands, which MW (indicated in kDa on the right) corresponded to dimers, trimers, tetramers, and pentamers of αABD (2α, 3α, 4α, and 5α, correspondingly). The band corresponding to hexamers (6α) was slightly visible upon blot overexpose. Right: the Oligomer Intensity Index (OII, the ratio between the intensities of the oligomer “n” to “n-1”) was quantified for 3α, 4α, 5α, and 6α based on five independent experiments. The ranges of distribution of the OII are shown. (**F**) Western blotting of GFP-αCatTCS-C805/871 and GFP-TCS-αABD-C805/871 cells probed with αABD antibody. (**G**) Fluorescence microscopy of αCatKO-A431 cells expressing GFP-TCS-αABD-C805/871 stained for GFP (GFP, green) and for F-actin (act, red). The left micrograph shows a merged image (Scale bar, 25 μm). Separate staining of the representative region (in a dashed box) is zoomed on the right (Scale bar, 15 μm). Note that the αABD is concentrated in tiny clusters along some actin-enriched structures. (**H**) Western blotting of cells expressing GFP-αCatTCS-C805/871 (αCat) and GFP-TCS-αABD-C805/871 (αABD) after permeabilization and thrombin treatment (as in E). Note that at least 8 adducts (marked as in E) could be detected in case of GFP-TCS-αABD-C805/871. The ranges of OII for each of the adduct are shown on the right (n= 5).

### α-Catenin oligomerization is adhesion- and force-independent

Aforementioned results showed that α-catenin binding to F-actin generates linear oligomers of CCC, which we refer to as CCC/actin strands. Does this process require cadherin *trans* interactions? To answer this question, we cultured cells in low calcium media that abolished cadherin *trans* interactions [42]. Surprisingly, this manipulation was able to neither decrease the levels of the adducts (Fig. 4A, lane LowCa), nor to change the adducts’ pattern found after thrombin cleavage (Fig. 4B). As another way to abolish E-cadherin *trans* interactions, we used the adhesion-blocking antibody SHE78-7. Lane SHE in figure 4A shows that this antibody also produced no effects on α-catenin oligomerization. LnB added for 20 min prior to cross-linking was sufficient to greatly reduce the amounts of α-catenin oligomers detected either in low calcium media or in the presence of SHE78-7 antibody (Fig. 4A, lanes LowCa+LnB and SHE+LnB). The oligomers reappeared 15 min after LnB washout (Fig. 4A, lane LowCa-postLnB). These observations provided strong evidence that cadherin *trans* interactions were not required for CCC strand formation.

**Figure 4.**
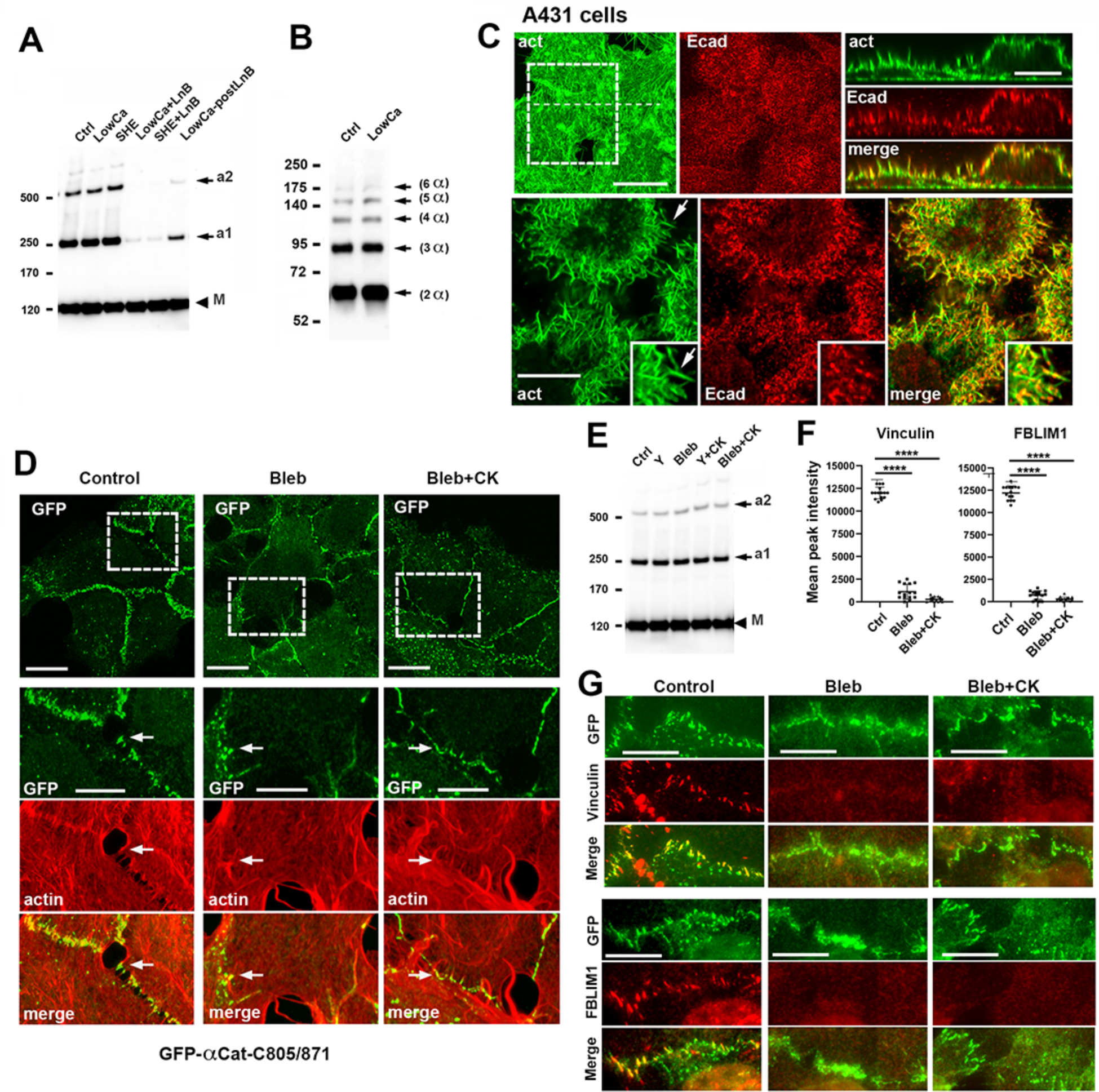
The α-catenin oligomers are adhesion- and myosin II-independent. (**A**) Western blot probed for GFP of GFP-αCatTCS-C805/871-expressing cells cross-linked after culturing in standard media (Ctrl), in low calcium media for 3 h (LowCa), in presence of function blocking E-cadherin antibody (SHE); in low calcium media or with SHE antibody in combination with LnB (LowCa+LnB, SHE+LnB), and as in lane LowCa+LnB but 15 min after LnB removal (LowCa-postLnB). Note that LnB addition dramatically reduced the level of the oligomers, which rapidly reappeared upon LnB washout (the bands marked as in Fig. 1E). (**B**) Western blot probed for αABD of GFP-αCatTCS-C805/871 cells in control (Ctrl) and Low calcium media (LowCa). The cells were permeabilized and treated with thrombin (as in Fig. 3E). Note that αABD oligomer pattern is the same. (**C**) Projections of all x-y optical slices of wt A431 cells stained before permeabilization for E-cadherin ectodomain (Ecad, red) and after permeabilization for F-actin (actin, green). Bar, 20 µm. The zoomed area (dashed box) of a single optical z slice passed through a middle of the cell is presented at the bottom. Bar, 10 µm. The zoomed area shown by the arrow is further magnified in the inserts. The optical z-cross-sections along the white dashed lines are shown on the right. Bar, 10 µm. Note that both, the isolated slice or the cross-section, showed clear co-localization between surface-exposed E-cadherin and F-actin. (**D**) Projections of all x-y slices of GFP-αCat-C805/871 cells stained for GFP (GFP, green) and for F-actin (actin, red). Only green channel is shown. Bar, 20 µm. The areas in dashed boxes are zoomed and shown in individual staining on the bottom. The arrow marks one of many actin-enriched AJs on each microphotograph. Bar, 10 µm. (**E**) Western blot probed for GFP of total lysates of GFP-αCatTCS-C805/871-expressing cells cross-linked after culturing in standard media (Ctrl), in presence of Y-27632 (Y), blebbistatin (Bleb), and in CK666 in combination with Y-27632 (Y+CK) or blebbistatin (Bleb+CK). (**F** and **G**) The parallel cultures of cells shown in D were stained for GFP and for tension-dependent proteins, vinculin or FBLIM1, and analyzed by immunofluorescent microscopy. (**F**) Quantification of peak intensities of vinculin and FBLIM1 in AJs. For quantification, the mean intensities of 30 AJs were determined. Statistical significance was calculated using a two tailed Student’s t test. ****, P < 0.0001. (**G**) Vinculin and FBLIM1 staining of the cells used for quantifications present in F. Only isolated cell-cell contacts are shown. Bar, 10 μm.

To visualize the CCC clusters on the cell plasma membrane in low calcium media, we stained the cells for E-cadherin ectodomain (before permeabilization) and for F-actin (after permeabilization). High resolution confocal microscopy of the cells cultured overnight in low calcium clearly showed numerous tiny E-cadherin clusters aligned with actin filaments (Fig. 4C). These clusters were especially abundant on the actin-rich plasma membrane protrusions (see insets in Fig. 4C). A parallel staining of these cells for β-catenin produced only weak background staining, confirming that the detected E-cadherin was localized to the plasma membrane. Furthermore, this background β-catenin staining showed little co-localization with actin filaments as judged by visual inspection (Fig. S1A) or by the values of Pearson’s correlation coefficient (PCC, Fig. S1B).

Our observation that the interactions of CCC with actin filaments are adhesion-independent suggested that αABD binding to actin is insensitive to the tensional stress on E-cadherin adhesions. To further validate this point, we treated the cells with the potent anti-actomyosin drugs, Rho-associated protein kinase (ROCK) inhibitor, Y-27632, or the myosin II inhibitor, blebbistatin (10 μM and 20 μM, correspondingly, for 1 hour prior to cross-linking). In order to remove the forces caused by the ARP2/3-facilitated actin flow [43, 44], these drugs were combined with an ARP2/3 inhibitor, CK666 (100 μM). In accordance with previous studies [30–33, 45], the drugs resulted in a dramatic rearrangement of AJs, from their typical radial appearance in control cells to more linear and dispersed in the treated cells, but they were unable to abolish AJ association with the actin cytoskeleton (Fig. 4D). Accordingly, neither of these drugs nor their combinations were able to change α-catenin oligomerization in our cross-linking assay (Fig. 4E). Staining for two actin-associated proteins, vinculin and FBLIM1, recruitment of which into AJs had been shown to be force-dependent [46, 47], confirmed that the drugs did release the tensional stress from the AJs (Fig. 4 F,G). Collectively, our results showed that αABD generates the actin-bound CCC strands independently of both actomyosin tension and *trans*-interactions of cadherin ectodomains.

### No other CCC proteins are needed for aABD to produce CCC/actin strands

To determine whether other known activities of CCC in addition to αABD-actin binding are needed to generate CCC/actin strands, we used a fusion protein EcΔ-GFP-α506C805/871. This chimera contains E-cadherin extracellular and transmembrane domains and an α-catenin region (aa 506-906) encompassing the M3 domain, a linker, and αABD (Fig. 5A). As demonstrated previously [48], this chimera produced actin-associated AJs in αCatKO-A431 cells (Fig. 5B). Our cross-linking assay also showed that this chimera, similar to the full-length α-catenin, formed actin-bound oligomers in both the control and in low calcium cultures (Fig. 5C). To test the role of *trans* E-clustering, we constructed a mutant, c/t-EcΔ-GFP-α506C805/871, which lacked both known *trans* dimer interfaces of E-cadherin (for X-dimers and strand-swap dimers) as well as its *cis* dimer interface (Fig. 5A). As expected, this mutant was unable to rescue cell-cell adhesion of the αCatKO-A431 cells or to produce AJs (Fig. 5D). Despite this, the cross-linking assay showed that both chimeras, the control one and the adhesion-incompetent, generated approximately the same levels of the actin-based oligomers (Fig. 5F).

**Figure 5.**
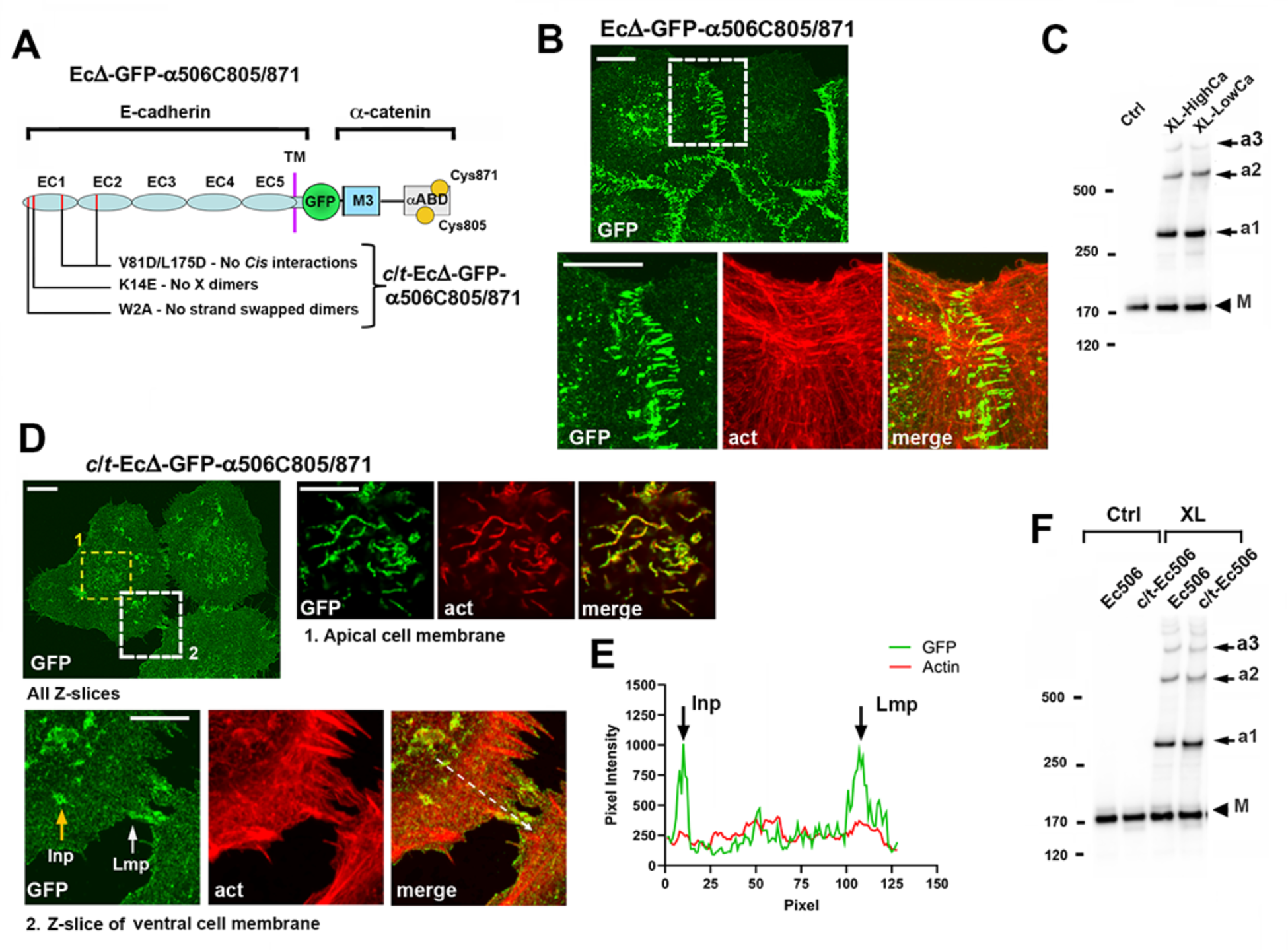
β-Catenin and p120 and are not needed to cluster E-cadherin through αABD. (**A**) Schematic representation of the E-cadherin-α-catenin chimera, Ec1′-GFP-α506C805/871, which consists of an intact N-terminal portion of E-cadherin (its ectodomain and transmembrane domain) fused through GFP with a C-terminal portion of α-catenin starting from M3 subdomain. Another chimera, c/t-Ec1′-GFP-α506C805/871, is the same as above but incorporates point mutations inactivating all known inter-ectodomain interactions: one *cis* and two *trans*, strand swapped and X dimerization. (**B**) Projections of all x-y optical slices of αCatKO-A431 cells expressing Ec1′-GFP-α506C805/871 stained for GFP (GFP, green) and for F-actin (actin, red). Only green channel is shown for low magnification. Bar, 10 µm. The dashed box area is zoomed on the bottom and shown as separate staining. Note that the chimera forms AJs associated with the actin cytoskeleton. Bar, 10 µm. (**C**) Western blot probed for GFP of of cells shown in B and collected from control culture (Ctrl) and from cross-linked cultures in standard media (XL-HighCa) or in low calcium media for 3 h (XL-LowCa). (**D**) Projections of all x-y slices of c/t-Ec1′-GFP-α506C805/871 cells stained for GFP (GFP, green) and for F-actin (actin, red). Only green channel is shown for low magnification. Bar, 10 µm. The dashed boxed areas are presented in both colors. Projections of five optical z slices encompassing the apical membrane marked by yellow box (#1) are zoomed on the right. Only a single optical slice passing through a ventral cell membrane (marked by a white box, #2) is shown on the bottom. Bars, 10 µm. Note that the apical ruffles, invadopodia-like structures (Inp, orange arrow), and cell-cell contacting lamellipodia (Lmp, white arrow) are especially enriched with the chimera clusters. (**E**) The line scan performed along the dashed line shown in D. Note that no specific actin enrichment is observed in places rich for the chimera clusters. (**F**) Western blot probed for GFP of cells expressing Ec1′-GFP-α506C805/871 (Ec506) and its adhesion-incompetent mutant (c/t-Ec506), which were obtained from control cultures (Ctrl) and after cross-linking (XL).

Confocal microscopy showed that the adhesion-incompetent chimera formed numerous actin-associated clusters. The clusters were especially abundant along the ruffles on the apical cell membrane (see zoomed area of the apical membrane in Fig. 5D) as well as on two types of actin-rich structures formed on the ventral cell membrane. One such structures was invadopodia (or their precursors), which are widespread in tumor cells [49]. The second type of structures was lamellipodia, especially often those that contact the neighboring cells. The intensity profile along the cell basal membrane showed that these two structures exhibited about five fold more GFP fluorescence than the surrounding cell membrane. By contrast, the level of F-actin in these sites was close to that in other areas of the basal membrane (Fig. 5E). The latter observation suggested that the clusters of the adhesion-incompetent chimera were driven by αABD oligomerization on actin filaments of particular actin cortex.

### CCC/actin strands are generated on myosin-1c rich actin cortex

Our previous experiments had been unable to detect clustering of the E-cadherin mutant WK-EcGFP, which, similar to c/t-Ecι1-GFP-α506C805/871, is defective for *trans* dimerization [50]. Our new findings prompted us to reinvestigate this issue because the high fluorescence of the non-clustered pool of this mutant could impede detection of small clusters by widefield microscopy, we had previously used. Indeed, despite a strong overall cytosolic and membranous fluorescence of the WK-EcGFP expressing cells (Fig. 6A, All Z-slices), confocal and deconvolution images of their apical and ventral plasma membranes did show that the mutant formed clusters. The clusters were concentrated along the same cortical domains as the clusters of the adhesion-incompetent chimera, c/t-Ecι1-GFP-α506C805/871, at ruffles, invadopodia, and lamellipodia (Fig. 6A, B). The plasma membrane localization of these clusters was confirmed by the staining of non-permeabilized cells using E-cadherin ectodomain mAb, HECD1 (Fig. S2).

**Figure 6.**
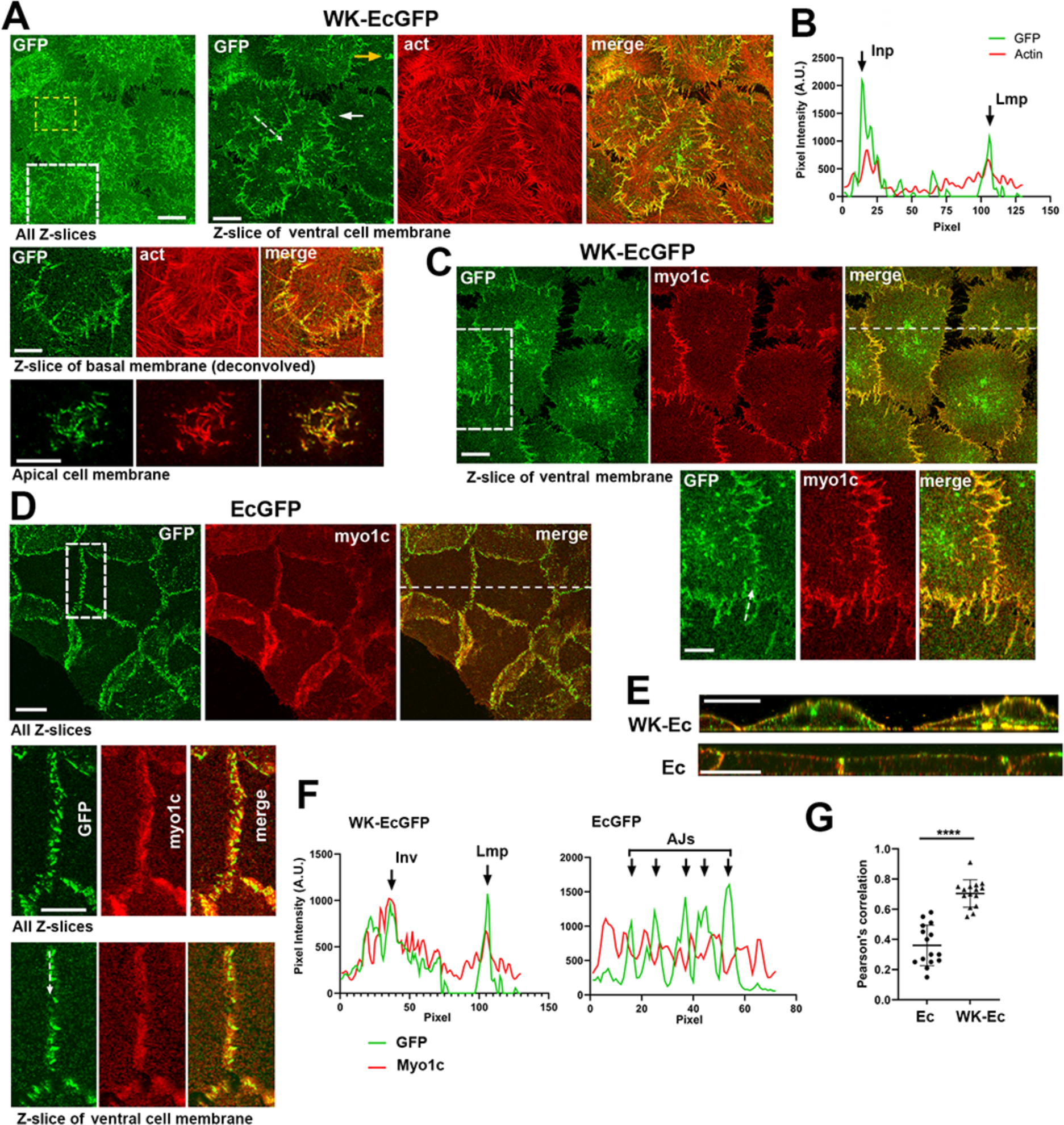
Adhesion-incompetent E-cadherin mutant interacts with a specific pool of actin filaments. (**A**) Representative A431(EP)-KO cells expressing WK-EcGFP mutant stained for GFP (GFP, green) and for F-actin (act, red). Left: projections of all optical z slices show a broad localization of the mutant obscuring its plasma membrane organization. Bar, 10 µm. Right: a single optical z slice of the ventral cell membrane taken from the left image. A white dashed box shown on the left image is deconvolved and zoomed in the second row (z-slice of ventral membrane, deconvolved). Projection of five optical z slices (spanning 1 µm of the apical cell region) of a yellow box shown on upper image is zoomed on the bottom. Bars, 5 µm. Note that the confocal slices show that WK-EcGFP is enriched in the apical raffles, in the invadopodia (yellow arrow), and cell-cell contacting lamellipodia (white arrow) of the ventral membrane. (**B**) The line scan performed along the dashed line shown in A. Note high concentration of the mutant in invadopodia (Inp) and in lamellipodia (Lmp). (**C**) WK-EcGFP cells were imaged for GFP (GFP, green) and myosin-1c (myo1c, red). Only a single optical z slice containing the ventral cell membrane is shown. Bar, 10 μm. The boxed area is zoomed on the bottom. Bar, 5 μm. The optical XZ cross section along the dashed line is shown in E. Note, an extensive co-distribution of the mutant and myosin-1c. (**D**) Projections of all z slices of the EcGFP-expressing cells stained for GFP (GFP, green) and myosin 1c (myo1c, red). Bar, 15 μm. The boxed area is zoomed on the bottom and given in form of all z projections (All z slices) or only as a single slice containing the ventral membrane (Z-slice of ventral membrane). Bar, 10 μm. (**F**) The line graphs of fluorescence intensities of GFP (green) and myosin-1c (red) along the white dashed lines shown in zoomed images in C and D (A.U., arbitrary units). Note that myosin-1c co-localizes with the WK-EcGFP in invadopodia (Inp) and in lamellipodia (Lmp) but is separated from EcGFP in AJs. (**G**) Average Pearson’s correlation coefficient between GFP and myoin-1c at the cell-cell contact areas in WK-EcGFP- and EcGFP-expressing cells. Correlations were calculated from 15 areas taken from 5 representative images of the single z slices of the basal cell membrane. A two tailed Student’s t test was used. ****, P < 0.0001.

The next question, therefore, was whether any actin-associated protein specifically marked the actin cortex that drove the WK-EcGFP clustering. To get an idea about possible candidates, we compared published WK-EcGFP and EcGFP proteomes [50]. Despite their high similarities, the proteome of WK-EcGFP showed some increase of myosin-1c, which, in fact, became its most abundant actin-binding protein. We therefore stained WK-EcGFP-expressing cells for myosin-1c. Remarkably, confocal microscopy showed that myosin-1c was concentrated in the same cortex domains as WK-EcGFP clusters (Figs. 6C and S3). Furthermore, close inspection and intensity profiles of cell-cell contact lamellipodia of these cells indicated significant colocalization of both proteins in these regions (Fig. 6C and F). Both proteins were also concentrated in the apical membrane ruffles (Fig. S3A). Similar association with myosin 1c was detected in the c/t-Ecι1-GFP-α506C805/871 expressing cells (Fig. S3B). A very different relationship was found between myosin-1c and intact E-cadherin. As in other epithelial cells [51], EcGFP-expressing cells exhibited myosin-1c in the basolateral cortex (Fig. 6D, E). However, the AJs, while located in the same basolateral domain, showed only slight or even no increase of myosine-1c staining compared to nearby areas (Fig. 6D, F). The differences in relationship between myosin-1c and functional and non-adhesive CCC was clearly reflected by PCC: its average value was relatively high (0.72) for WK-EcGFP, but only 0.36 for EcGFP (Fig. 6G). These results suggest that the *trans* cadherin interactions triggered segregation of CCC from the myosin-1c rich actin cortex.

### CCC/actin strands integrate into AJs by redundant mechanisms

In addition to the actin-binding unit of αABD, this domain encompasses another evolutionary conserved region, the αABD tail. The tail (aa 872-906) is disordered in all available αABD models and is not needed for αABD binding to F-actin [8, 9, 24, 52]. It had been recently identified that the tail provides an actin-bundling activity to αABD [25]. This activity of αABD could potentially be important for cadherin clusters. To test this, we expressed an α-catenin mutant with truncated tail, GFP-αCatΔ892-C805/871, in αCatKO-A431 cells (Fig. 7A, B). In agreement with the in vitro binding data, our cross-linking approach showed no impact of this deletion on formation of the CCC strands (Fig. 7B, lane Δ892). However, contrary to our expectations, the tail truncation also produced no gross abnormalities in cell-cell adhesion, as judged by visual inspection of the cells, nor in association of AJs with actin bundles (Fig 7C, column Δ882, see also below).

**Figure 7.**
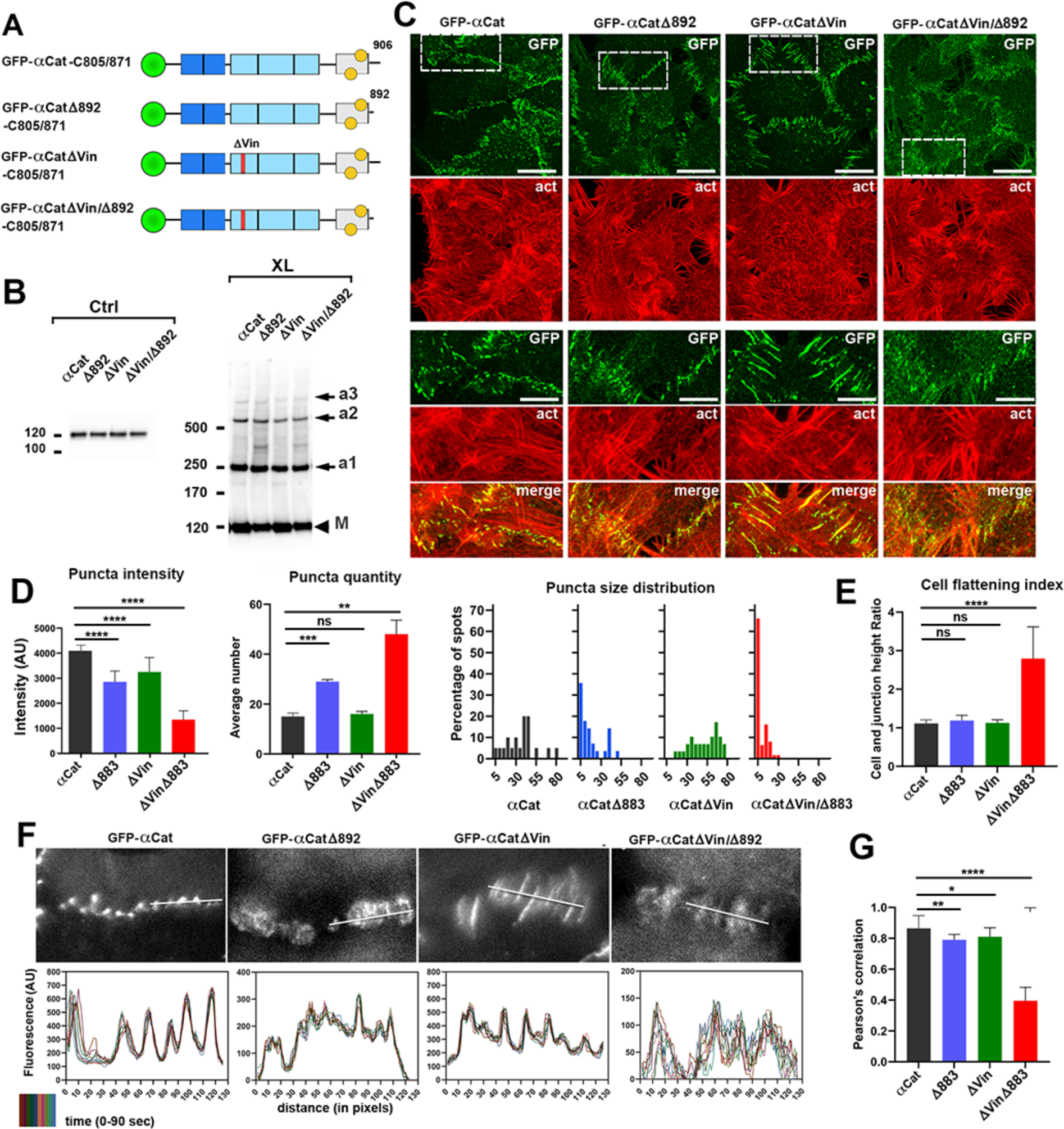
Impact of αABD tail deletion on cell-cell adhesion. (**A**) Diagram of αABD tail mutants of α-catenin. Only a set of the mutants bearing cysteine replacements are shown. The color code of the α-catenin subdomains is the same as in Figure 1A. Note that the αABD tail truncation (1′892) was introduced into α-catenin and into its vinculin uncoupled mutant (1′Vin). Another set of the mutants, with larger truncation (1′883) and without cysteine replacements is not shown. (**B**) The cells expressing the set of the mutants shown in A were probed in Western blotting in control culture (Ctrl) or after cross-linking (XL). Note that the cross-linking generated the same adducts in all four cultures. (**C**) Confocal fluorescence microscopy of the cells expressing the set of the mutants with cysteine replacements (the mutant names are indicated on the top). The cells were stained for GFP and F-actin. Projections of all optical z slices are shown. Bars, 15 μm. The boxed regions are zoomed on the bottom. Bars, 5 μm. (**D**) Morphometric analyses of AJs. The AJs were determined as bright puncta in cell-cell contacts with areas 1 pixel or more. Note that the cells expressing α-catenin double mutant (1′Vin1′883) were strikingly different from other three clones by all parameters: fluorescence intensity of puncta (the value of brightest pixel), by quantity of puncta (per 10×10 μm of the contact), and by their size (number of pixels in the puncta). (**E**) Cell flattening index (the ratio between the height of the cells to the height of the apical junctions (see Fig. S4 for detail) that reflects the general strength of cell-cell adhesion. (**F**) Time-lapse microscopy of the cells expressing α-catenin mutants taken at 10 sec temporal resolution. An isolated frame taken from the representative movies (Videos M1-M4) of each cell clone are shown in the top row. The representative line scans along the white lines of 10 consecutive frames (that span 1.5 min) of each of the movies were combined and differently colorized. The combined scans show that the cells expressing the double mutant form highly unstable AJs. (**G**) Cell-cell contact instability was also determined as the average PCC between consecutive frames of the movies presented in F. Statistical significance for all graphs was calculated using a two tailed Student’s t test: ns, non-significant; *, P < 0.05; **, P<0.01; ***, P< 0.001; ****, P < 0.0001.

One of the possibilities we pursued was that vinculin, which also can bundle actin, compensated for the lack of the αABD tail-mediated bundling activity. We tested this idea using an additional mutation, ΔVin, that inactivates the vinculin-binding site of α-catenin [24]. As expected, the new mutants GFP-αCatβVin-C805/871 and GFP-αCatΔVinΔ892-C805/871 (Fig. 7A) showed no abnormalities in αABD oligomerization in our cross-linking assay (Fig. 7B). Also, in agreement with the previous data [24, 53], the GFP-αCatβVin-C805/871 mutant, in which the ΔVin mutation was combined with the intact αABD, induced elongation of AJs along the radial cell axis but resulted in no clear impacts on the general cell phenotype (Fig. 7C, column ΔVin). However, in combination with the tail truncation, mutation of the vinculin-binding site produced a dramatic effect on cell-cell adhesion (see below). These results were fully replicated using another set of mutants (GFP-αCat, GFP-αCatβVin, GFP-αCatΔ883, and GFP-αCat-βVin/β883), which lacked cysteine substitutions and bore an alternative truncation of the αABD tail. They confirmed that the dramatic effects of the combinatorial mutation on AJs were not based on the accidental adverse effects of the particular mutant.

The dual α-catenin mutation, 1′Vin/1′883 (or 1′Vin/1′892), led to a significant disintegration of AJs (Fig. 7C). Instead of characteristic micrometer-sized adhesive structures present in other cells, the cells with this mutant formed numerous submicron puncta. This visual observation agreed with our morphometric analysis that included quantification of the fluorescence intensities of junctional structures, their numbers and sizes (Fig. 7D). It showed that compared to the GFP-αCat-expressing cells, the average number of α-catenin puncta increased by a factor of 3 in the dual mutant cells. Concomitantly, their fluorescence intensities dropped by ∼ 70 %. The most striking difference was the size of the puncta: the pool of smallest junctions (with the area of 5 pixels or less), which was just ∼5% in the control cells, increased to more than 60% in the cells expressing the dual mutant. By all these parameters, the cells expressing α-catenin with just the 1′Vin mutation behaved similarly to the control cells. The tail truncation alone induced the intermediate phenotype (Fig. 7D).

A dysfunction of AJs containing the combinatorial mutant resulted in clear defects in the shape of cells in single-layered cell cultures (Fig. S4). The ratio between the height of the cell and the height of their apical junctions, that showed the flattening of the cells, was close to one for the cells expressing the wild type or single α-catenin mutants, indicating that they were relatively flat, but it was close to three in the cells expressing the double α-catenin mutant (Fig. 7E). This difference showed that the cell-cell adhesion of the latter cells is not sufficient for their flattening. Time-lapse microscopy taken with 10 sec temporary resolution also showed that the junctions of the dual mutant cells differed from other by their instability (Videos 1-4). To quantify this, we generated line scans of the representative cell-cell contacts from the movies (Fig. 7F) and calculated PCC between individual frames (Fig. 7G). The results showed that the AJs of control cells and the cells expressing only ΔVin or only Δ883 (or Δ892) mutants remained stationary for 1-2 minutes. By contrast, the AJs with the combinatorial 1′Vin/1′883 (or 1′Vin/1′892) mutant changed their position and appearance even between two consecutive frames.

As we showed above, matured AJs and adhesion-incompetent CCC/actin strands interacted with different pools of actin cortex that could be distinguished by staining for myosin-1c. We, therefore, stained the cells expressing different α-catenin mutants for myosin 1c (Fig. 8). These data clearly showed that the intact α-catenin, as well as the mutant bearing a single 1′Vin mutation, were separated from myosin-1c equally well, as validated by relatively low (∼ 0.35) values of PCC (Fig. 8C). The combinatorial mutant showed the best colocalization, with PCC average 0.6, close to that for WK-EcGFP (see Fig. 6G).

**Figure 8.**
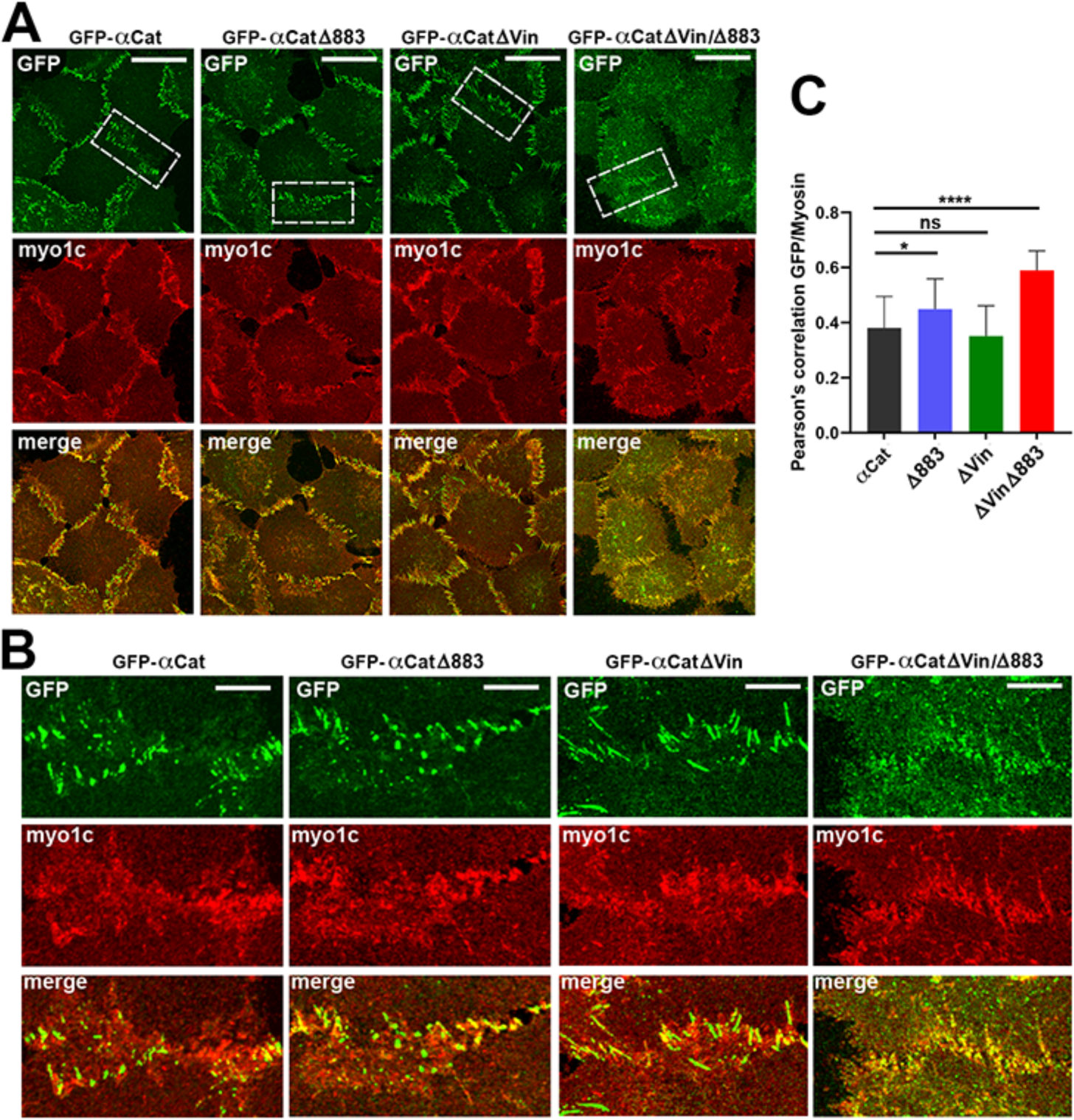
Relationship between AJs and myosin-1c in cells with α-catenin mutants. (**A**) Confocal fluorescence microscopy of the cells expressing the set of the mutants with αABD truncation (the names of the mutants are indicated on the top). The cells were stained for GFP (GFP) and myosin-1c (mio1c). Projections of all optical z slices are shown. Bars, 25 μm. (**B**) The zoomed representative cell-cell contacts indicated by dashed line boxes in A. Bars, 7 μm. Note that most AJs in control cells and the cells expressing α-catenin with a single mutation do not show high correlation with myo1c. By contrast, the puncta incorporated α-catenin with double mutation are concentrated in myo1c-rich areas. (**C**) GFP-myosin-1c co-localization was also determined as the average PCC. Statistical significance was determined by a two tailed Student’s t test: ns, non-significant, P>0.05; *, P < 0.05; ****, P < 0.0001.

## Discussion

It is largely accepted that direct binding of α-catenin to F-actin through αABD plays two critical roles in cell-cell adhesion. The first beeing stabilization of cadherin clusters that is needed to upregulate avidity of the adhesive AJ interface [19, 20, 54, 55]. The second is sensing the actomyosin contraction forces that control AJ maturation, and the recruitment of vinculin, in particular [56–60]. In addition, αABD binding to actin is proposed to play a third critical function – it could drive cadherin clustering [24]. This idea has been suggested by a remarkable cooperativity of αABD binding to actin. The underlying mechanism of this behavior is partially explained by a cryo-EM model of the actin-binding interface of αABD: it utilizes the residue, Val870, from the αABD attached to the neighboring site on the filament [8, 9, 25]. Such dependency on the neighbors results in oligomerization of αABD along the filaments. To demonstrate this process in cells, we engineered an α-catenin mutant enabling specific detection of such linear αABD oligomers using targeted cross-linking approach. Experiments with the designed mutant clearly show that αABD binding to actin does result in αABD oligomerization exactly as suggested by the cryo-EM blueprint. The resulting oligomers consist of up to 6 α-catenin molecules, while the most abundant oligomeric species is a pentamer. Since α-catenin of the oligomers is incorporated into CCC, this oligomerization should place CCC along the filament forming actin-bound linear arrays, which we define as CCC/actin strands.

Unexpectedly, our results show that these CCC/actin strands are adhesion independent. Accordingly, low calcium media or a cadherin function-blocking antibody produce no effects on the strand formation. The strands generated by the E-cadherin-α-catenin chimera are also unchanged upon inactivation of both *trans* and *cis* binding sites of the chimera ectodomain. Such non-adhesive CCC/actin strands are detected by confocal microscopy as small actin-associated cadherin clusters in low calcium media or in cells expressing adhesion incompetent mutants of E-cadherin or E-cadherin-α-catenin chimera. Furthermore, the strands are also independent from forces caused by motor activity of myosin II and by Arp2/3-based actin polymerization: the drugs inhibiting these proteins are unable to reduce CCC/actin strand formation. The non-adhesive CCC/actin strands we detected might correspond to the previously described cadherin *cis* oligomers observed by various fluorescence techniques on the contact-free membrane [16, 61, 62]. In contrast to the fluorescence techniques, which identify cadherin oligomers with no structural details, the targeted cross-linking approach detects specific supramolecular structures.

The non-adhesive CCC strands are predominantly observed on the membrane protrusions generated by a branched actin network, such as ruffles, lamellipodia or invadopodia. The buildup of CCC/actin strands on the plasma membrane of cell protrusions, where they should be in rapid equilibrium with monomeric CCC, explains a large body of data identifying cell protrusions as key sites for AJ formation [63–70]. Noteworthy, we show that the CCC/actin strands forming on the protrusive structures are juxtaposed with myosin-1c, a member of the class I myosin superfamily. The latter proteins are known for their role in specific arrangement of actin filaments in cell protrusions and lateral domains of epithelial cells [51, 71]. It is tempting to speculate that myosin-1c contributes to the specific organization of the actin cortex that promotes formation of the αABD-actin strong bonds. One of the possibilities is that this motor protein produces tension across actin filaments that was shown to enhance αABD binding to actin [9].

In standard cultures of epithelial cells, the majority of CCC is recruited into cell-cell contacts. Previous studies have shown that a key mechanism of this recruitment is the entrapment of E-cadherin in the contacts by formation of *trans* E-clusters, the ordered crystalline-like 2D lattice, which spontaneously assembles by cooperation of *cis* and *trans* interactions of the cadherin ectodomain [1, 2, 13–15, 73]. The CCC/actin strands and E-clusters are fully compatible since the inter-protomer distance in the strands (∼ 6 nm) is close to that (∼ 7 nm) in E-clusters [7–9]. The most likely scenario therefore is that in cell-cell contacts the CCC/actin strands integrate into E-clusters forming composite oligomers, which we refer to as *trans* CCC/actin clusters (Fig. 9). Importantly, the incorporation of just one CCC/actin strand into a *trans* E-cluster, which itself is too unstable to play a role in adhesion [19], should increase the overall binding energy of the cluster thereby increasing its adhesive capacity. A possibility for an intermix of actin-bound and actin-uncoupled CCC in the same cluster has been shown by efficient incorporation of the tail-deleted E-cadherin mutant into the AJs build by endogenous E-cadherin [15]. Incomplete saturation of AJs with CCC strands is also indicated by the observation reported here that ∼ 50% of E-cadherin in AJs is soluble in Triton X100 buffers. Altogether, an obvious hypothesis is that AJs at each given moment consist of numerous *trans* CCC clusters incorporating various numbers of CCC strands. Such organization corresponds well to the immuno-EM images of cadherin clusters formed between the cells and the cadherin-coated substrate [20]. Many of those clusters have been observed as straight lines of three to six gold nanoparticles (∼ 6 nm each) that fit the characteristics of CCC strands. More complex clusters could arise from a growth of these strands through E-clustering and from the addition of new strands.

**Figure 9.**
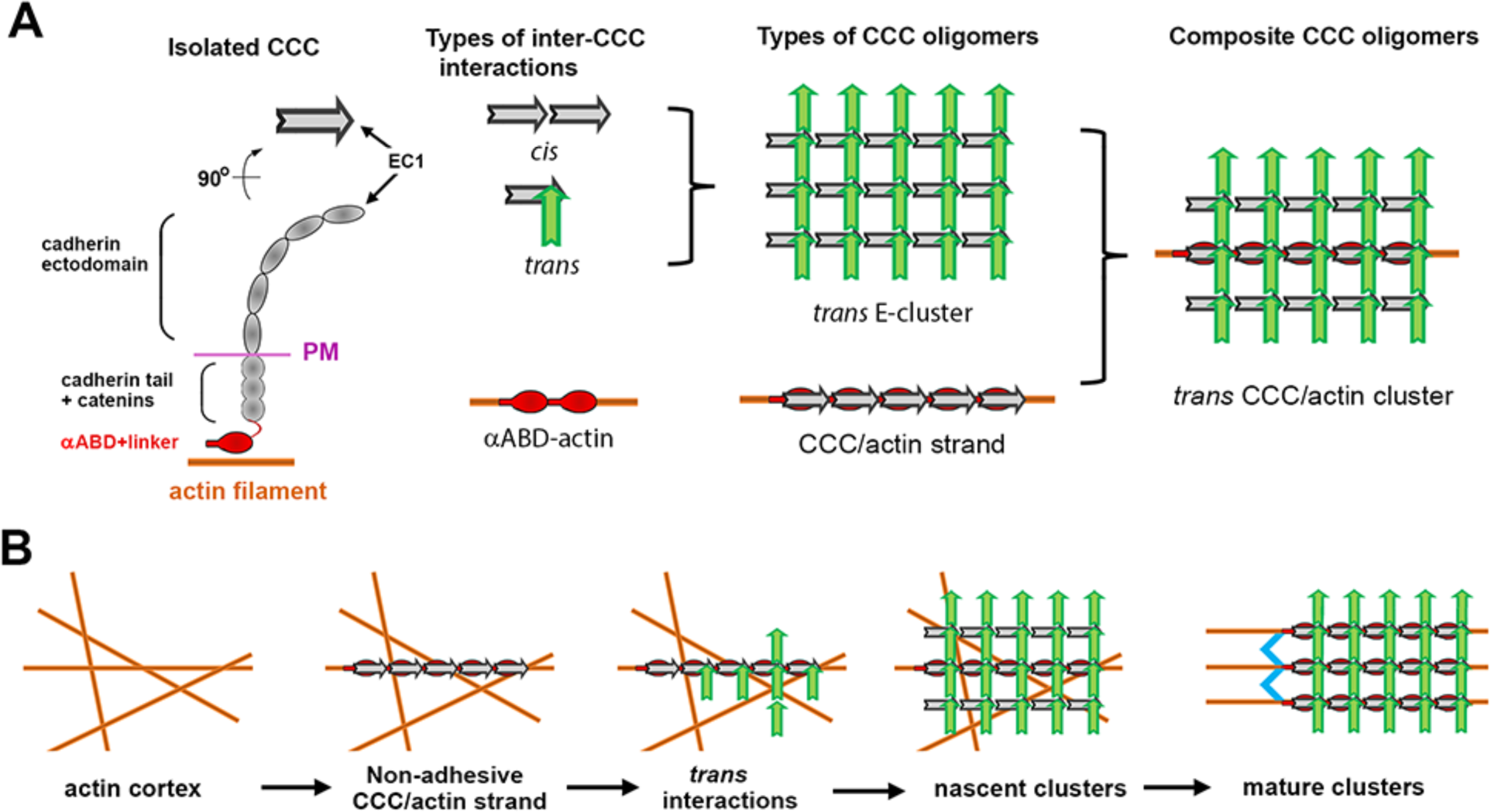
Modeling of E-cadherin clusters and their interactions with the actin cytoskeleton. (**A**) **Types of CCC oligomers**. Simplified representation of an isolated CCC (*Isolated CCC*). The side view shows its extracellular region (*ectodomain*), depicted by five ellipsoids corresponding to cadherin extracellular domains (the amino-terminal EC1 domain is marked), the plasma membrane region (PM), and the intracellular cadherin tail in complex with catenins (cadherin tail+catenins). The actin-binding domain of α-catenin separated from the rest of CCC by a flexible linker is shown in red (αABD+ linker). The top view projection of the ectodomain is shown even in more simplified form. It is depicted by an arrow, which pointed end corresponds to the EC1 domain. Three types of interactions between CCCs are known (*Types of inter-CCC interactions*): two modes of binding are known for the ectodomain, *cis*-binding (*cis*) and *trans*-binding (*trans*, the ectodomains colored in grey and green belong to lower and upper cells correspondingly). Note that *trans*-interacting ectodomains are perpendicular one another. Inside the cells CCCs interact in *cis* through the αABD-actin interactions (*αABD-actin*). The inter-CCC interactions may spontaneously produce two types of oligomers (Types of CCC oligomers). The ectodomain *cis* and *trans* interactions form *trans* E-clusters in cell-cell interface [reviewed in 42, 72]. The αABD-actin interaction forms CCC/actin strands (this study). Finally, both oligomerization processes could be intermixed forming composite CCC oligomers (*trans* CCC/actin cluster). (**B**) **Tentative multistep mechanism of cadherin clustering**. The short-lived CCC/actin strands are continuously generated on the specialized actin cortex. As soon as the strands appear in the cell-cell interface, the strands nucleate assembly of the *trans* E-clusters thereby forming nascent actin-associated cadherin clusters. To function in adhesion, these clusters should be further modified by rearrangement of the associated actin filaments using F-actin cross-linking activities (depicted in blue).

Formation of *trans*-CCC clusters appears to be only an initial step in AJ assembly. We identify here two complementary mechanisms that provide the clusters with the additional strength needed to establish a functional adhesive site. The first one is based on vinculin, the α-catenin-related protein that binds to α-catenin under force [58, 59, 73]. Vinculin is known for its role in reorganization of the branched actin network into bundles at the sites of nascent cell-substrate adhesions [74, 75]. The second one is based on αABD tail. This highly conserved unstructured 30 aa-long portion of αABD is not involved in αABD binding to F-actin [8, 24, 52]. One of the reported activities of this region is also a bundling of the αABD-associated filaments by a yet unknown mechanism [25]. These two bundling activities may redundantly contribute to the local rearrangement of actin filaments that results in segregation of maturated AJs from a myosin-1c-rich cortex. Accordingly, only minor defects in AJs could be detected in cells expressing α-catenin with targeted inactivation of its vinculin binding site or its αABD tail. Like control cells with the intact α-catenin, cells expressing either of these mutants still assemble AJs, which are associated with the actin bundles and are well isolated from myosin-1c. By contrast, the cells expressing an α-catenin mutant combining these two mutations cannot form AJs. While this dual mutant still produces CCC strands, the strands are dispersed along numerous submicron puncta that are retained in the myosin-1c-containing cortex. The puncta, in contrast to AJs, undergo rapid, in the order of seconds, continuous reorganization. The importance of actin filament reorganization from branched networks to parallel filament bundles has been previously shown for maturation of focal adhesions [76–79].

In summary, our findings show that the onset of cadherin adhesions is the integration of two CCC oligomerization processes that occurs in cell-cell contact interface. The first one produces actin-bound linear CCC/actin strands, while the second one generates 2D *trans* E-clusters. While connected to actin filaments, the resulting *trans*-CCC/actin clusters are inactive in adhesion. To get activated, the clusters need to rearrange their actin filaments by filament bundling activities associated with CCC. This late step of cluster assembly contributes to formation of a symbiotic CCC-actin filament matrix, known as an AJ plaque, which facilitates high reassembly dynamics of the individual clusters and their interconnection with the actomyosin cytoskeleton [80]. Molecular insight into the late step is imperative to fully elucidate the mechanisms of cell-cell adhesion and its defects in various human diseases.

## Materials and Methods

### Plasmids

The plasmids (all in pRcCMV) encoding GFP-tagged human αE-catenin (denoted GFP-αCat), its mutant GFP-αCat-1′Vin, in which its vinculin binding site is inactivated by five amino acid substitutions to alanine (R329A, R330A, L347A, L348A, and Y351A), as well as GFP-tagged human E-cadherin (EcGFP) and its mutant WK-EcGFP, in which its *trans* binding interface is inactivated by two point mutations, W2A and K14E, have been reported [24, 50]. The E-cadherin-α-catenin chimera EcΔ-GFP-α506 was also previously described [48]. PCR-based mutagenesis of these plasmids was used for constructing all other mutants described here. The general maps of all mutants are presented in Figs. 1, 3, and 7. All plasmid inserts were verified by sequencing.

### Cell culture and transfection

The original A431 cells, A431(EP)-ko cells expressing EcGFP and WK-EcGFP, and α-catenin deficient αCatKO-A431 cells have been previously described [50]. Other cells were obtained using stable transfection with the corresponding plasmids of the A431(EP)-ko or αCatKO-A431cells as indicated. The cells were grown in DMEM supplemented with 10% FBS and were transfected using Lipofectamine 2000 (Invitrogen) according to the company protocol. After selection of the Geneticin-resistant cells (0.5 mg/ml), the cells were sorted for transgene expression by FACS, and only moderate-expressing cells were used. At least three clones were selected for each construct, and all were tested in most of the assays. The levels and sizes of the recombinant proteins in the obtained clones were analyzed by Western blotting. All clones of cells expressing a particular transgene exhibited the same phenotype. Representative data for one of three clones is presented.

For the Ca^2+^-switch assay, cells were cultivated in a low Ca2+ medium (20 µM Ca^2+^) for the indicated time. The drugs were added for 30 min at the following concentrations: Latrunculin B (1 μM), Y-27632 (10 μM), blebbistatin (20 μM), CK666 (100 μM).

### Cross-linking, SDS-PAGE, immunoprecipitation, and thrombin cleavage

For cross-linking, 2-day-old confluent cultures grown in 24-well tissue culture dishes were washed in phosphate-buffered saline (PBS) supplemented with 0.5 mM CaCl_2_ (PBS-C) and cross-linked by incubation for 5-min on ice with ice-cold PBS containing 40 μM of cysteine-specific cross-linker BMB (ThermoFisher). The reaction was stopped by washing the cells with PBS with dithiothreitol (1 mM). The cells were then lysed in SDS sample buffer and the adducts were analyzed by Western blotting as previously described [81]. In brief, the total lysates were separated on precast 3-8% Tris-acetate gels (Invitrogen), which are ideal for the separation of large MW proteins. A molecular weight protein marker (10–460 kDa; HiMark #LC5699, Invitrogen) was used for MW analyzes.

The coimmunoprecipitation assays have been performed as described previously [81]. In brief, confluent A-431 cells expressing indicated recombinant proteins grown on 10 cm plates were extracted with 1.5 ml of IP-buffer (50 mM Tris-HCl, pH 7.4, 150 mM NaCl, 0.5 mM AEBSF, 2 mM EDTA, and 1% Triton X-100). The insoluble material was removed by centrifugation and the lysates were subjected to immunoprecipitation either by subsequent incubations with anti-Ecad antibody SHE78-7 (4 μg/ml) and protein A-beads or by incubation with GFP-trap beads (Chromotek). After incubation, the beads were washed four times in IP-buffer, boiled in 30 μl of SDS-sample buffer, and loaded on SDS-PAGE gels. For non-crosslinked samples the precast 4-15% Tris-Glycine Bio-Rad gels were used. Restriction grade thrombin cleavage kit (Novagen, 69671-3) was used for thrombin cleavage as indicated by manufactory protocol using thrombin dilution 1:250 and 30 min incubation time at RT in the provided thrombin cleavage buffer. For cell extraction (or permeabilization before thrombin cleavage), the 1% triton X100 in the cytoskeleton preservation buffer (100 mM PIPES, pH 6.9; 1 mM MgCl_2_; 1 mM EGTA) was used.

### Immunofluorescence microscopy

For immunofluorescence, cells were grown for 2 days on glass coverslips or imaging glass-bottom dishes (P35G-1.5; MatTek) and were fixed with 3% formaldehyde (5 min) and then permeabilized with 1% Triton X-100, as described previously [82]. Wide-field images (Figs. 1D, 2A, 3B, G, 4G) were taken using an Eclipse 80i Nikon microscope (Plan Apo 100×/1.40 objective lens) and a digital camera (CoolSNAP EZ; Photometrics, Tucson, AZ). The confocal images were taken using a Nikon AXR laser scanning microscope equipped with a Plan Apo 60x×/1.45 objective lens. Immediately before imaging, the dishes were filled with 90% glycerol. The images were then processed using Nikon’s NIS-Elements software.

For immunostaining the following antibodies were used: mouse anti-E-cadherin mAb clones SHE78-7 and HECD1 (Takara, M126 and M106), mouse anti-vinculin (Sigma, V9264), chicken anti-GFP (Novus, NB100-1614), rabbit anti-FBLIM1 (Novus, NBP2-57310), anti-β-catenin (Invitrogen, PA5-16762), anti-myosin 1c and anti-α-catenin (Abcam, ab194828 and ab51032), anti-αABD (Cell Signaling, 36611). All secondary antibodies were produced in Donkey (Jackson Immunoresearch Laboratories). Alexa-Fluor-555–phalloidin and Latrunculin A were purchased from Invitrogen.

### Live-cell imaging

The live cell imaging experiments were performed essentially as described previously [15, 82] using a halogen light source. In brief, cells were imaged (in IP-buffer, Fig 2E, or in L-15 media with 10% FBS, Fig. 7) by an Eclipse Ti-E microscope (Nikon, Melville, NY) at RT or 37°C controlled with Nikon’s NIS-Elements software. The microscope was equipped with an incubator chamber, a CoolSNAP HQ2 camera (Photometrics) and a Plan Apo VC 100x/1.40 lens. The 2×2 binning mode was used in all live-imaging experiments. At this microscope setting, the pixel size was 128 nm. All images were saved as Tiff files and processed using ImageJ software (National Institutes of Health).

### Data processing

All images were processed and analyzed using Nikon’s NIS-Elements ver. 5.02. For line scan analysis, the Element’s in-built line profile function was used to draw a 1pix wide line across the junctions. For peak intensity measurement, the highest intensity on the Y-axis was recorded. A minimum of 15 independent junctions were scanned from five different images. For the size and the number of puncta analyses, the Binary function of NIS-Elements AR ver. 5.02 was used. In brief, the Element’s in-built threshold function was selected to identify bright circular objects of moderate to heavy clustering. All spots with a diameter starting from 1px and above were chosen with a background threshold of 30 arbitrary units that correspond to fluorescence intensity outside the cell-cell contacts. The number of remaining puncta (per 10×10 μm) and their respective sizes were exported to excel and were used for calculations using GraphPad Prism 10.2.0. For Pearson’s correlation quantification, the images were processed with limited background reduction and denoising function of NIS-Element 5.02 and then Element’s Pearson function was used to measure the correlation between green and red fluorescence of the selected cell-cell contact areas (10×10 μm) of the confocal images or between the consecutive frames of the movies. Representative 15 areas taken from 5 images, and 14 different consecutive frames from 3 movies were analyzed. The charts and error bars were plotted using GraphPad Prism version 10.2.0. Statistical significance was analyzed using student’s two-tailed t test for two groups and ANOVA analysis for more groups. A p value that was less than 0.05 was considered statistically significant.

For measuring cell and junction height, the NIS-Element’s slices view was used. A new document was created from the slices view and the built-in Annotations and Measurements function was used to draw two two-point lines, one of which from the base of the cell to its most apical junction and another one from the base to the top of the nucleus to measure the junctional height and the cell height, correspondingly. The ratio was calculated using cell height and junctional height of the cell. A minimum of 20 different heights were measured from 5 different confocal images. The charts and error bars were plotted using GraphPad Prism version 10.2.0.

## Acknowledgments

We thank Drs. T. Svitkina (University of Pennsylvania), J.W. Mitchell and B.J. Mitchell (Northwestern University) for valuable comments and suggestions. The study was also stimulated by fruitful discussions of α-catenin structure with Drs. B. Honig and L. Shapiro (all Columbia University, New York). Sequencing, flow cytometry and confocal microscopy were performed at the Northwestern University Genetic, Flow Cytometry, and Advanced Microscopy Centers. The authors declare no competing financial interests. The work was supported by National Institute of Health Grant AR070166 and GM148571 (to S.M.T.).

## Figure legends

**Figure S1.**
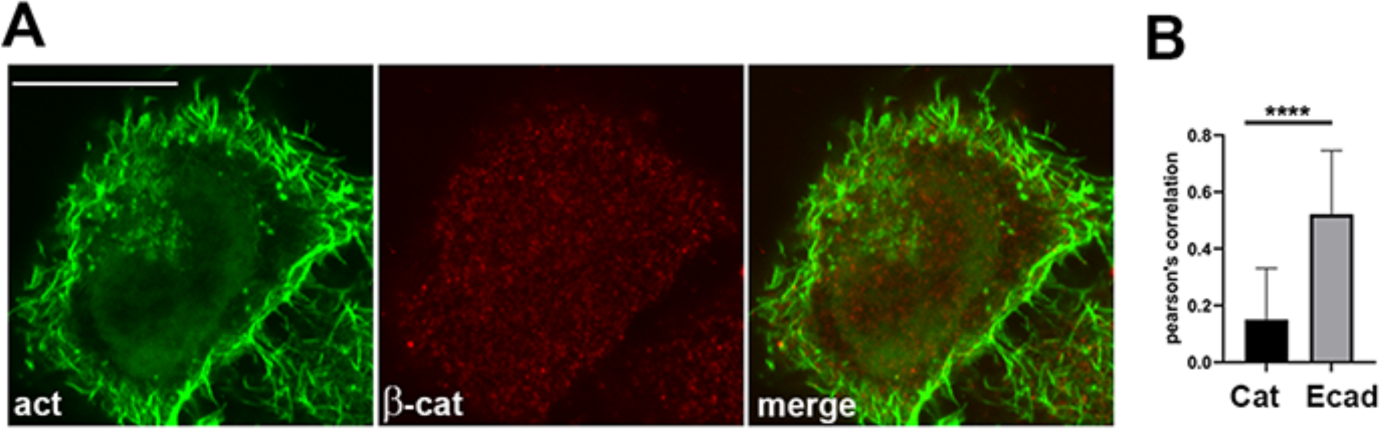
The α-catenin oligomers are adhesion-independent. (**A**) A single optical z slice passed through a middle of the wt A431 cell stained before permeabilization for β-catenin (βCat, red) and after permeabilization for F-actin (actin, green). Bar, 20 µm. Note that the antibody produces only background staining with no clear co-localization with F-actin. (**B**) Average Pearson’s correlation coefficient (PCC) of β-catenin/F-actin (shown in A) and E-cadherin/F-actin (shown in Fig. 4C). Correlations were calculated from 15 areas taken from 5 representative single z slices passed through middle of the cells. A two tailed Student’s t test was used. ****, P < 0.0001.

**Figure S2.**
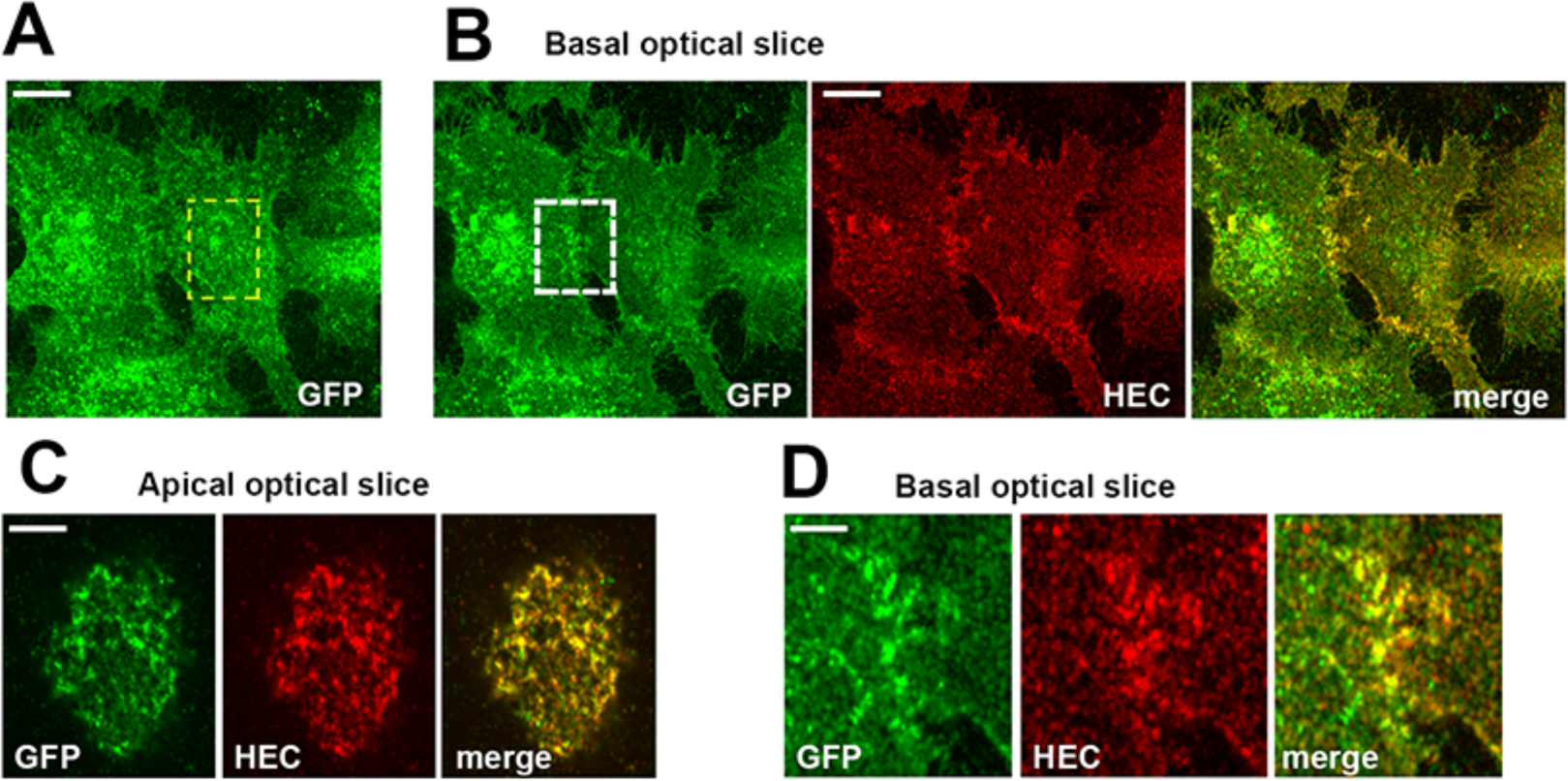
The clusters of the adhesion-incompetent WK-EcGFP mutant are exposed on the cell surface. Representative A431(EP)-KO cells expressing the adhesion incompetent WK-EcGFP mutant stained for E-cadherin ectodomain by mAb HECD1 (HEC, red) before permeabilization and after permeabilization for GFP (GFP, green) (**A**) Projections of all x-y optical slices. Only GFP staining is shown. Bar, 15 µm. (**B**) A single optical z slice of the basal plasma membrane taken from the stack shown in A. (**C**) Zoomed portion of the apical plasma membrane demarcated by yellow dashed line in A. Projection of five optical z slices (spanning 1 µm of the apical cell region) is shown. Note that the WK-EcGFP clusters reside on the apical raffles. Bar, 5 µm (**D**) Zoomed portion of the basal plasma membrane demarcated by white dashed box in B. Note, that the mutant forms numerous tiny plasma membrane-located clusters. Bar, 5 µm.

**Figure S3.**
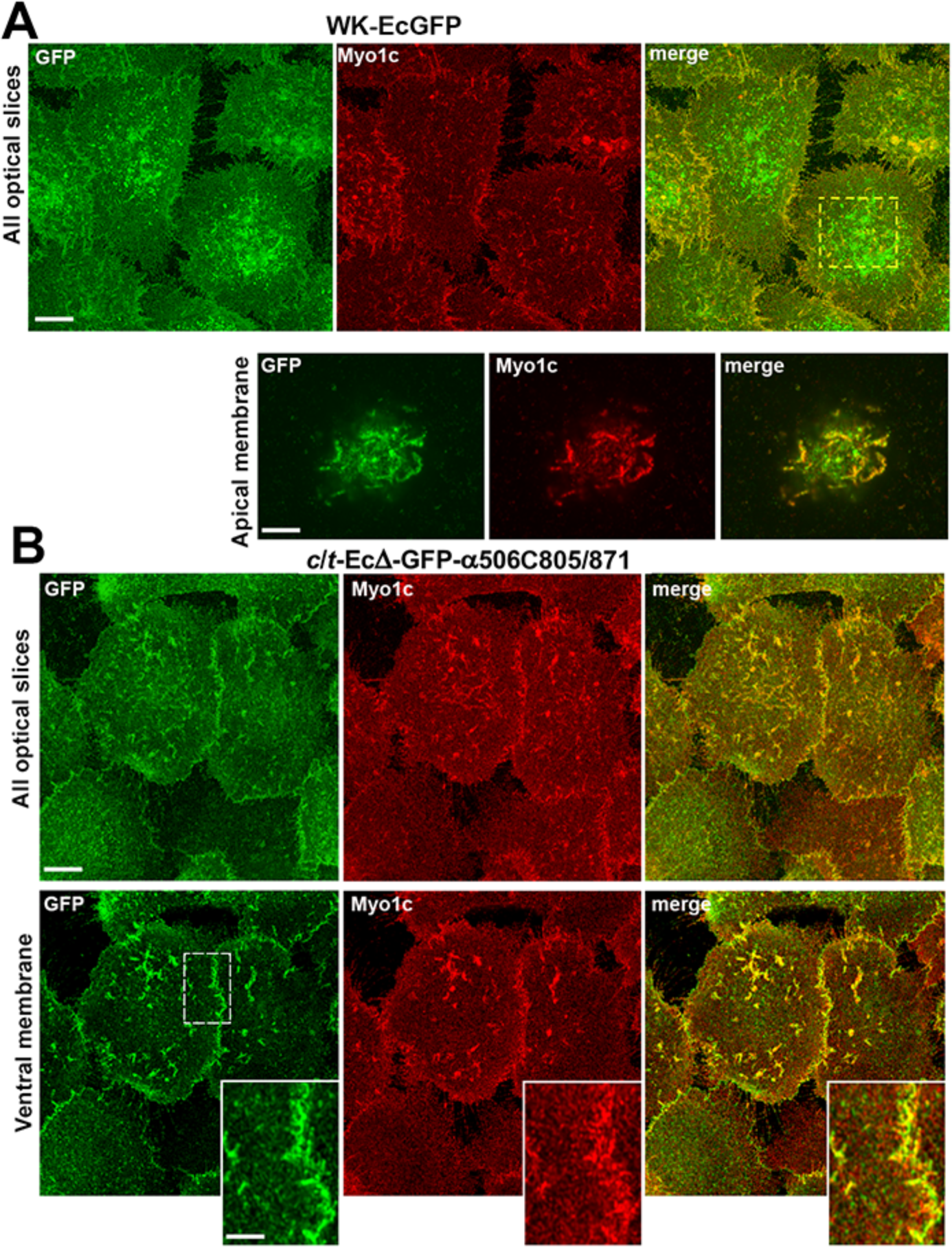
The non-adhesive clusters associate with myosin-1c enriched actin cortex. (**A**) Projections of all optical z slices of the cells expressing WK-EcGFP shown in Fig. 6C (All optical slices). Note a broad localization of the mutant (GFP). Bar, 10 µm. Projections of only 5 apical z slices of the boxed area is zoomed on the bottom (apical membrane). Bar, 5 μm. Note that clusters of the mutant are colocalized with myosin-1c. (**B**) The α-catenin-deficient A431 cell expressing the adhesion-incompetent chimers c/t-EcΔ-GFP-α506C805/871. The cells were stained for GFP (GFP, green) and myosin-1c (Myo1c, red). Projections of all optical z slices (top) and only the slice containing the ventral membrane (bottom) are shown. Bar, 10 µm. Note, the chimera clusters and myosin-1c are localized on the same lamellipodia. The area indicated by dashed line is zoomed in the insets. Bar, 5 μm.

**Figure S4.**
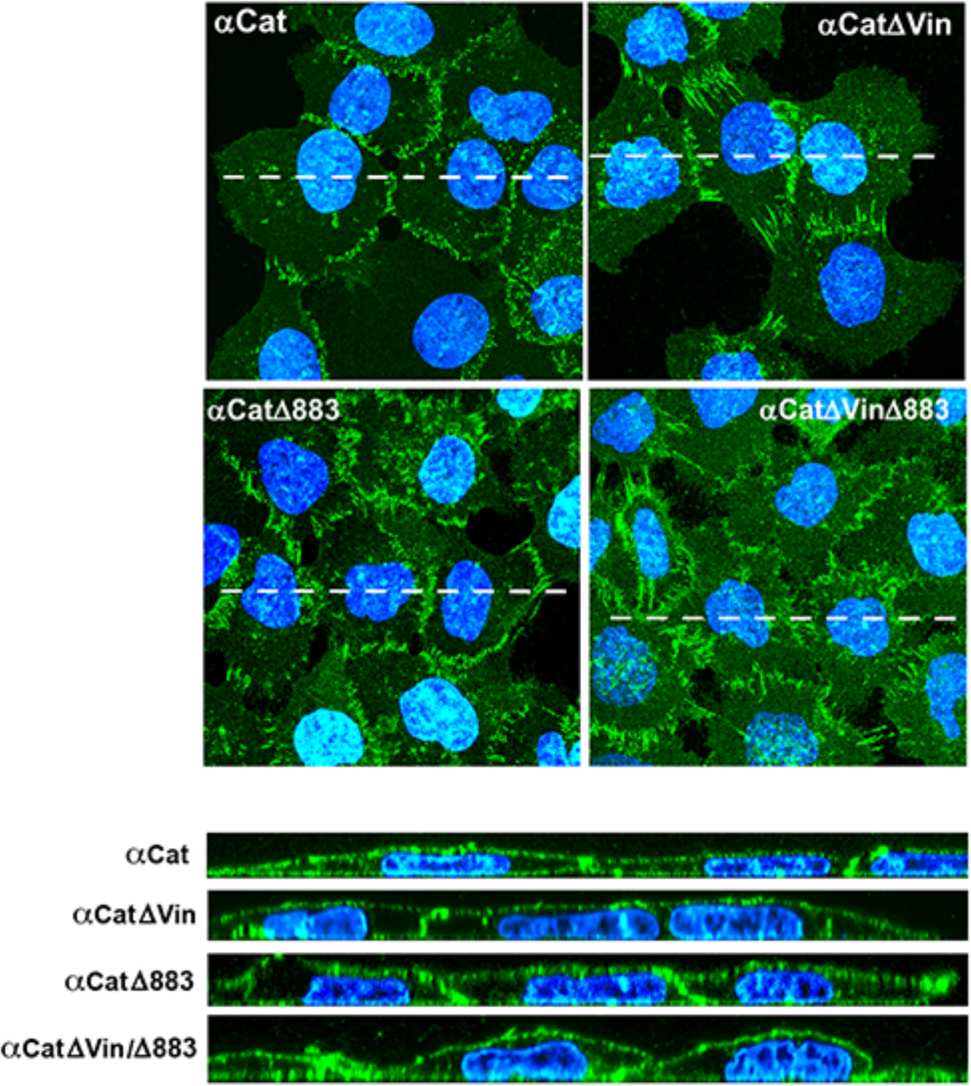
The cells expressing the dual α-catenin mutant are not flattened. Confocal fluorescence microscopy of the cells expressing the set of the mutants with αABD truncation (the names of the mutants are indicated). The cells were stained for GFP (green) and DAPI (blue) to visualize nuclei. Projections of all z slices are shown. Bars, 25 μm. The optical XZ cross sections along the dashed lines are shown on the bottom. Note that the apical plasma membrane of the double mutant expressing cells are curved, so the top of the nucleus is higher than the apical junctions.

**Video S1. Dynamics of the AJs in αCat-GFP-expressing in αCatKO-A431 cells**. The cells were imaged in control media and images were acquired at 10 sec intervals.

**Video S2. Dynamics of the AJs in αCat-GFP1′883-expressing in αCatKO-A431 cells**. The cells were imaged in control media and images were acquired at 10 sec intervals.

**Video S3. Dynamics of the AJs in αCat-GFP1′Vin-expressing in αCatKO-A431 cells**. The cells were imaged in control media and images were acquired at 10 sec intervals.

**Video S4. Dynamics of the AJs in αCat-GFP1′Vin1′883-expressing in αCatKO-A431 cells**. The cells were imaged in control media and images were acquired at 10 sec intervals. Note that AJs of these cells exhibit extremely high motion suggesting that AJs of these cells were unable to reorganize their cytoskeleton.

## Notes

### Competing Interest Statement

The authors have declared no competing interest.

